# *De novo* mutations in KIF1A-associated neuronal disorder (KAND) dominant-negatively inhibit motor activity and axonal transport of synaptic vesicle precursors

**DOI:** 10.1101/2021.07.22.453457

**Authors:** Yuzu Anazawa, Tomoki Kita, Rei Iguchi, Kumiko Hayashi, Shinsuke Niwa

## Abstract

KIF1A is a kinesin superfamily molecular motor that transports synaptic vesicle precursors in axons. Mutations in *Kif1a* lead to a group of neuronal diseases called KIF1A-associated neuronal disorder (KAND). KIF1A forms a homodimer and KAND mutations are mostly *de novo* and autosomal dominant; however, it is not known whether the function of wild-type KIF1A is inhibited by disease-associated KIF1A when they are dimerized. No reliable *in vivo* model systems to analyze the molecular and cellular biology of KAND caused by loss of function mutations have been developed; therefore, here, we established *Caenorhabditis elegans* models for KAND using CRISPR/cas9 technology and analyzed defects in axonal transport. In the *C. elegans* models, heterozygotes and homozygotes exhibited reduced axonal transport phenotypes. Suppressor screening using the disease model worm identified a mutation that recovers the motor activity of disease-associated human KIF1A. In addition, we developed *in vitro* assays to analyze the motility of single heterodimers composed of wild-type KIF1A and disease-associated KIF1A. Disease-associated KIF1A significantly inhibited the motility of wild-type KIF1A when heterodimers were formed. These data indicate the molecular mechanism underlying the dominant nature of *de novo* KAND mutations.

**Significance Statement:** KIF1A is a molecular motor that transports synaptic vesicle precursors in axons. Recent studies have identified many *KIF1A* mutations in congenital neuropathy patients; however, the molecular mechanism of pathogenesis remains largely elusive. This study established loss of function models for KIF1A-associated neuronal disorder (KAND) in *Caenorhabditis elegans* to analyze the molecular and cell biology of the disease *in vivo*. Genetic screening using the disease model could find a mutation that recovers the motor activity of disease-associated KIF1A. This study also established *in vitro* single-molecule assays to quantitatively analyze the effect of KAND mutations when mutant KIF1A forms heterodimers with wild-type KIF1A. Our findings provide a foundation for future genetic screening and for drug screening to search for KAND treatments.

## Introduction

Neuronal function depends on intracellular transport system (1). Kinesin superfamily proteins (KIFs) and a cytoplasmic dynein are molecular motors for anterograde and retrograde transport, respectively (2, 3). Various membranous organelles and protein complexes in neurons are anterogradely transported by Kinesin-1, -2 and -3 family members (4-8). Neurons transmit information via synaptic vesicles that are accumulated to synapses in the axon (9). The constituents of synaptic vesicles are synthesized and assembled in the cell body and transported down axons to synapses by a mechanism called axonal transport. The transported organelle is called a synaptic vesicle precursor (7). Kinesin superfamily 1A (KIF1A), a Kinesin-3 family member, is an axonal transport motor for synaptic vesicle precursors (7, 10). KIF1A have a motor domain and a cargobinding tail domain (7). The motor domain, conserved among Kinesin superfamily members, has microtubule-dependent ATPase activity that drives movement on microtubules (11, 12). The tail domain of KIF1A is composed of a protein-binding stalk domain and a lipid binding Pleckstrin-homology (PH) domain (10, 13-15).

*Caenorhabditis elegans* (*C. elegans*) is a good model animal to study axonal transport (16-23). UNC-104 is a *C. elegans* orthologue of KIF1A (6, 24). Electron and light microscopy analyses have shown synapses as well as synaptic vesicles are mislocalized in *unc-104* mutants (6). The mechanism of axonal transport is well conserved between *C. elegans* and mammals and the expression of human *Kif1a* cDNA can rescue the phenotype of *unc-104* mutant worms (25).

Mutations in the motor domain of KIF1A cause congenital neuropathies (26, 27). More than 60 mutations have been found in the motor domain of KIF1A in neuropathy patients. Some cases are familial, but most are sporadic. For example, KIF1A(R11Q) was found in a spastic paraplegia patient who has autism spectrum disorder (ASD) and attention-deficit hyperactivity disorder (ADHD) (28). KIF1A(R254Q) mutation was found in Japanese spastic paraplegia patients with intellectual disability (29). KIF1A(R254) is a hot spot for a broad range of neuropathies. KIF1A(R254W) have been described in patients from other countries (27). These broad range of neuropathies, caused by KIF1A mutations, are called KIF1A-associated neuronal disorder (KAND) (27). Both dominant and recessive mutations are associated with KAND. Recent *in vitro* studies have shown that most of KAND mutations are loss of function. KIF1A(V8M) have defects in force generation (30). KIF1A(A255V) and KIF1A(R350G) have shorter run length (31). KIF1A(P305L) mutation reduces microtubule association rate of the motor (32). KIF1A(R169T) disrupted the microtubule-dependent ATPase activity of the motor domain(33). KIF1A(R254W) mutation reduces the velocity and run length of the motor protein (27). On the other hand, we have suggested that KIF1A(V8M), KIF1A(A255V) and KIF1A(R350G) mutations, all of them are familial, are gain of function (25). *In vitro* analysis using the full-length human KIF1A as well as worm models established by CRISPR/cas9 suggested these mutations disrupts the autoinhibitory mechanism and overactivate KIF1A, leading to increasing of the axonal transport of synaptic vesicle precursors.

While loss-of-function mutations have been intensively studied using *in vitro* assays, reliable models to study the neuronal cell biology of loss-of-function KAND mutations *in vivo* are awaited. Moreover, previous *in vitro* studies have mostly analyzed homodimers composed of disease-associated KIF1A (27, 32) because it was difficult to purify heterodimers composed of wild-type and disease-associated KIF1A(30). Activated KIF1A forms a homodimer to move on microtubules (34); therefore, wild-type KIF1A is very likely to dimerize with disease-associated KIF1A in patient neurons. However, properties of heterodimers composed of wild type KIF1A and disease-associated KIF1A remains largely unknown, and it is elusive whether *de novo* KAND mutations inhibit the function of wild-type KIF1A in a dominant negative fashion.

Here, we established worm models of *de novo* KAND mutations. Heterozygous worms, as well as homozygous worms, show synaptic deficiencies that are caused by axonal transport defects. Unbiased suppressor screening using the worm model identified a mutation that recovers the motor activity of disease-associated KIF1A. We also established an *in vitro* single molecule analysis system to measure the motility parameters of a single heterodimer composed of wild-type and disease-associated KIF1A. Heterodimers composed of wild-type KIF1A and disease-associated KIF1A showed reduced motility *in vitro*. Both *in vitro* and *in vivo* analysis suggest a dominant-negative nature of disease-associated mutations.

## Result

### *C. elegans* models of *de novo* KAND

To study molecular and cellular deficiencies caused by *de novo* disease-associated KIF1A mutations, we established *C. elegans* models for KAND using CRISPR/cas9 (35). *C. elegans unc-104* gene is an orthologue of human *Kif1a*. We introduced following mutations in *unc-104* gene: *unc-104(R9Q), unc-104(R251Q)* and *unc-104(P298L)* (Fig 1A and B and supplementary Fig S1A). These UNC-104 residues are conserved in human KIF1A and these mutations correspond to KIF1A(R11Q), KIF1A(R254Q) and KIF1A(P305L) mutations, respectively. All of them are causes of *de novo* and autosomal dominant KAND(28, 29, 32). Introduction of mutations was confirmed by digestion with restriction enzymes and Sanger sequencing (Fig S1A). Then, we observed the macroscopic phenotypes of disease model homozygous worms. As a control, a strong loss-of-function allele of unc-104, *unc-104(e1265)*, was included. *unc-104(e1265)* is described as *unc-104(lf)* in this paper. Homozygous worms showed strong uncoordinated (unc) phenotypes and did not move well on the culture plate (Fig 1B). To quantitively analyze the movement of worms, the number of body bends in a water drop was counted during minute (Fig S1B). We found *unc-104(R9Q), unc-104(R251Q), unc-104(P298L)* did not move well in the water as is the cause in *unc-104(lf)*. Moreover, the body size of homozygous worms was smaller than wild type (1.09 ± 0.09 mm, 0.66 ± 0.06 mm, 0.64 ± 0.07 mm, 0.74 ± 0.06 mm and 0.81 ± 0.14 mm, mean ± standard deviation in length, respectively in *wild type, unc-104(R9Q), unc-104(R251Q), unc-104(P298L)* and *unc-104(lf)*) (Fig 1C and S1C). These results collectively show that KIF1A(R11Q), KIF1A(R254Q) and KIF1A(P305L) mutations are loss of function.

**Figure 1.**
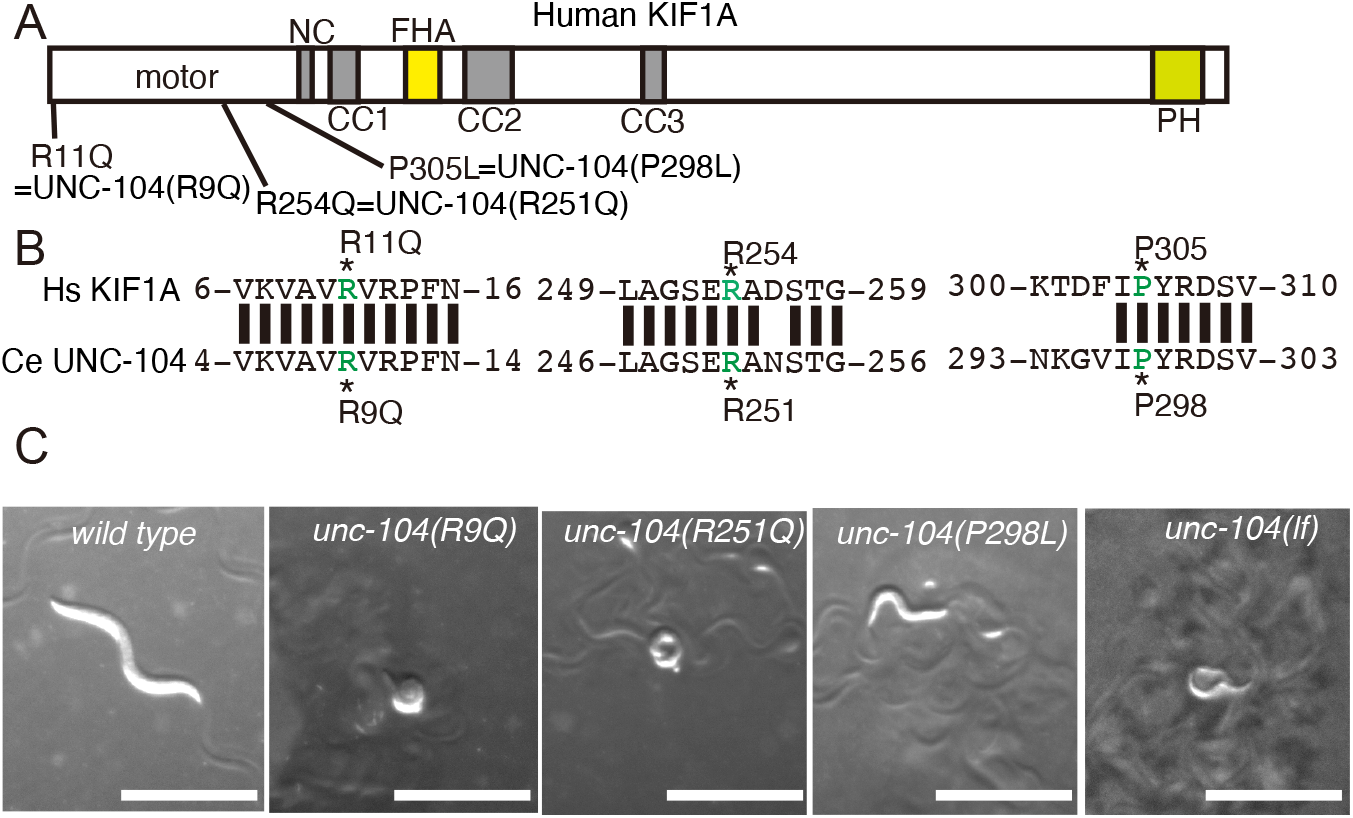
(A) Schematic drawing of the domain organization of KIF1A motor protein. NC, neck coiled-coil domain. CC1, Coiled-coil 1 domain. FHA, Forkhead-associated domain. CC2, Coiled-coil 2 domain. CC3, Coiled-coil 3 domain. PH, Pleckstrin-homology domain. The three KAND mutations and corresponding *C. elegans* UNC-104 mutations analyzed in this study are indicated. (B) Sequence comparison between human KIF1A and *C. elegans* UNC-104. (C) Macroscopic phenotypes of KAND model homozygotes at 1-day adults. Mutant worms are smaller than wild type worms and do not move well on the bacterial feeder. Bars, 1 mm. See also supplementary Figure S1.

### Synaptic vesicles are mislocalized in homozygotes

UNC-104 is an orthologue of human KIF1A and is a molecular motor that determines the localization of synaptic vesicles in *C. elegans*; therefore, we visualized synaptic vesicles in KAND model worms. The DA9 neuron in *C. elegans* is highly polarized and forms *en passant* synapses along the dorsal side of the axon (36) (Fig 2A). The characteristic morphology of DA9 neuron is suitable for analyzing axonal transport and synaptic localization (37). We expressed a synaptic vesicle marker GFP::RAB-3 in the DA9 neuron using the *itr-1* promoter to visualize DA9 synapses (Fig 2B). In KAND models, GFP::RAB-3 signals were reduced in the axon and strongly misaccumulated in the dendrite (Fig 2C,2D and supplementary Fig S2). Only a trace amount of GFP::RAB-3 signal was observed in the DA9 axon in KAND models.

**Figure 2.**
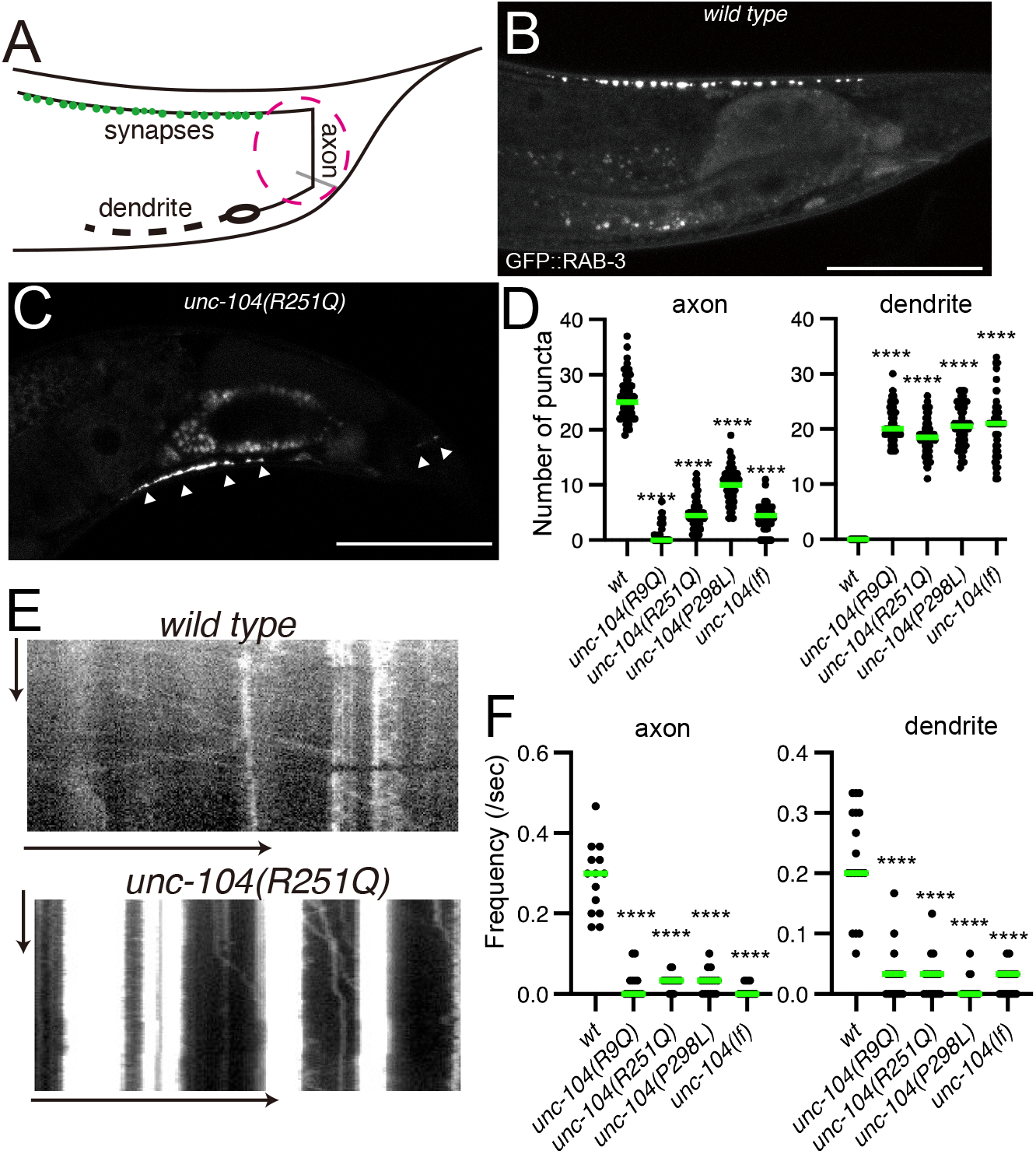
Synaptic vesicle localization in KAND model homozygote worms. (A) Schematic drawing show the morphology of DA9 neuron. Green dots along the axon show synaptic vesicle distribution. The red circle shows proximal axon. (B and C) Representative images showing the distribution of synaptic vesicles in the DA9 neuron in *wild type* (B) and *unc-104(R251Q)* (C). Synaptic vesicles are visualized by GFP::RAB-3. Arrowheads show mislocalization of synaptic vesicles in the dendrite and proximal axon. Bars, 50 *μ*m. (D) Dot plots showing the number of puncta in the dorsal axon (left panel) and ventral dendrite (right panel) of DA9. Green bars represent median value. Kruskal-Wallis test followed by Dunn’s multiple comparison test. N = 60 worms for each genotype. ****, Adjusted P value < 0.0001. (E) Representative kymographs in *wild type* (upper panel) and *unc-104(R251Q)* (lower panel). The axonal transport of synaptic vesicle precursors was visualized by GFP::RAB-3. The proximal axon shown in panel (A) was observed. Vertical and Horizontal bars show 10 seconds and 10 *μ*m, respectively. (F) Dot plots showing the frequency of anterograde axonal transport (left panel) and retrograde axonal transport (right panel). Green bars represent median value. Kruskal-Wallis test followed by Dunn’s multiple comparison test. N = 14 *wild type*, 14 *unc-104(R9Q)*, 18 *unc-104(R251Q)* and 16 *unc-104(P298L), unc-104(lf)* axons. ****, Adjusted P Value < 0.0001. See also supplementary Figure S2.

We then observed axonal transport of synaptic vesicle precursors in the proximal region of the DA9 axon (37) (Fig 2A, magenta circle). We used GFP::RAB-3 as a representative marker for axonal transport of synaptic vesicle precursors because previous studies have shown that GFP::RAB-3 co-migrates with other synaptic vesicle and pre-synaptic proteins in the axon and is, therefore, a good marker to visualize axonal transport (10, 23, 37). In the *wild type* worms, both anterograde and retrograde transport were observed in the axon (Fig 2E and F). In contrast, the frequency of both anterograde and retrograde events was significantly reduced in all three mutant strains (Fig 2E and F). In more than 70% mutant worms, no vesicular movement was detected in the 30 sec time window. Similar phenotypes are observed in loss of function allele of *unc-104*. These data indicate that axonal transport of synaptic vesicles is strongly reduced in *unc-104(R9Q), unc-104(R251Q)* and *unc-104(P298L)* strains.

### KAND mutations disrupt the motility of motor proteins *in vitro*

To study the effect of KAND mutations *in vitro*, we analyzed the motility of purified human KIF1A protein using total internal reflection fluorescence (TIRF) microscopy (38) (39). To directly study motility parameters, regulatory domains and cargo binding domains were removed (Fig 3A). The neck coiled-coil domain of mammalian KIF1A does not form stable dimers without cargo binding (40); therefore, we stabilized human KIF1A dimers using a leucine zipper domain as described previously (27) (32). A red fluorescent protein, mScarlet-I, was added to the C-terminal of the protein to observe the movement (Fig 3A). Resultant KIF1A homodimers [KIF1A(1-393)::LZ::mSca] were purified by Strep tag and gel filtration (Fig 3B and supplementary Fig S3A). All the recombinant proteins were recovered from the same fractions in the gel filtration. The recombinant protein was then used to analyze the motility of single KIF1A dimers on microtubules (Fig 3C-J). The motility of KIF1A(1-393)::LZ::mSca dimers was detected at 10 pM (Fig 3C). KIF1A(1-393)(R11Q)::LZ::mSca did not move at all on microtubules even at 100 pM (Fig 3D) while KIF1A(1-393)::LZ::mSca was saturated on microtubules under the same condition (Fig 3G). KIF1A(1–393)(R11Q)::LZ::mSca showed only one-dimensional Brownian motion on microtubules. KIF1A(1-393)(R254Q)::LZ::mSca moved on microtubules at 10 pM (Fig 3E). We observed a binding frequency of KIF1A(1-393)(R254Q)::LZ::mSca with microtubules is higher than KIF1A(1-393)::LZ::mSca (Fig 3I) but the average velocity was 50% lower and the average run length was 70 % shorter than for wild type (Fig 3H and J). The landing rate and run length of KIF1A(1–393)(P305L)::LZ::mSca was significantly lower than wild type (Fig 3I), consistent with previous studies (27, 32). Although affected parameters were different depending on the mutated residues, these data are consistent with the reduced axonal transport phenotypes observed in KAND model worms.

**Figure 3.**
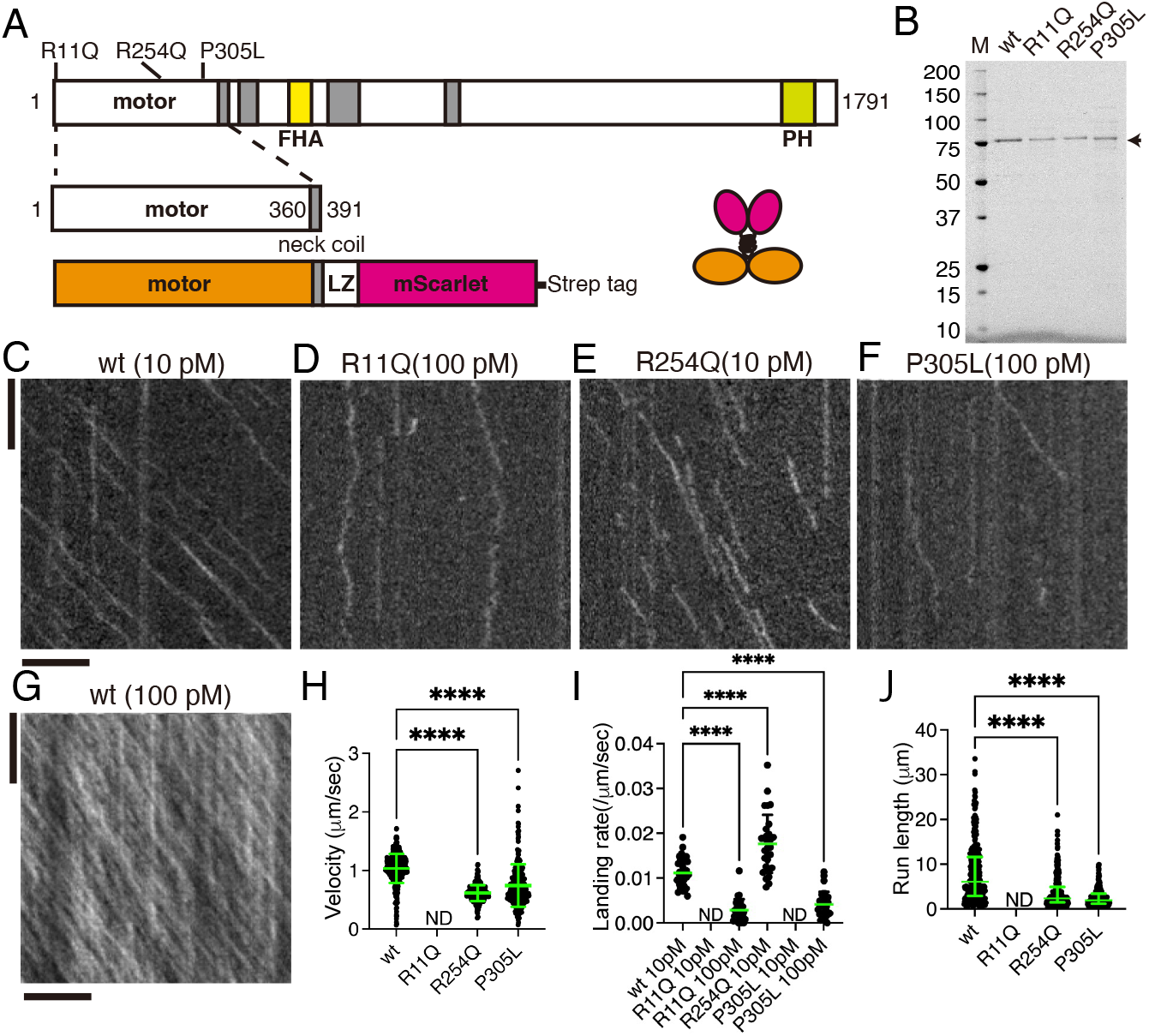
Single molecule behavior of disease-associated KIF1A mutants. (A) Schematic drawing of the domain organization of KIF1A motor protein and recombinant protein analyzed in Figure 3. (B) Purified KIF1A(1–393)::LZ::mScarlet and its mutants were separated by SDS-PAGE and detected by trichloroethanol staining. M represents a marker lane. Numbers on the left indicate molecular weight (kDa). Arrow indicates KIF1A(1–393)::LZ::mScarlet. (C-G) Representative kymographs showing the motility of 10 pM KIF1A(wt) (C), 100 pM KIF1A(R11Q) (D), 10 pM KIF1A(R254Q) (E), 100 pM KIF1A(P305L) and 100 pM KIF1A(wt) (G). Vertical and horizontal bars represent 5 sec and 5 *μ*m, respectively. (H) Dot plots showing the velocity of KIF1A. Each dot indicates one datum.. Green bars represent mean ± standard deviation (S.D.). Kruskal-Wallis test followed by Dunn’s multiple comparison test. n = 433 (wt), 325 (R254Q) and 498 (P305L). ****, Adjusted P Value < 0.0001. Note that no processive movement was detected for KIF1A(R11Q). (I) Dot plots showing the landing rate of KIF1A. The number of KIF1A that bound to microtubules was counted and adjusted by the time window and microtubule length. Each dot shows one datum. Green bars represent median value. Kruskal-Wallis test followed by Dunn’s multiple comparison test. n = 30 (10 pM wt), 28 (100 pM R11Q), 29 (10 pM R254Q) and 30 (100 pM P305L) movies. ****, Adjusted P Value < 0.0001. Compared with KIF1A(wt). Note that no landing event was detected in 10 pM KIF1A(R11Q) and KIF1A(P305L) experiments. (J) Dot plots showing the run length of KIF1A. Each dot shows one datum. Green bars represent median value and interquartile range. Kruskal-Wallis test followed by Dunn’s multiple comparison test. n = 312 (wt), 241 (R254Q) and 243 (P305L) homodimers. ****, Adjusted P Value < 0.0001. Note that all KIF1A motility events were included, including those that end when the motor reaches the end of an MT; thus, the reported run lengths are an underestimation of the motor’s processivity. See also supplementary Figure S3.

In contrast to well characterized KIF1A(P305L) mutation (32), properties of KIF1A(R11Q) and KIF1A(R254Q) mutations have not been analyzed well. Then, we compared the binding of these two mutants with microtubules in the presence of ADP and an AMP-PNP (ATP analogue) (Figs S3B-E). The binding of KIF1A(R254Q) with microtubules is comparable to KIF1A(wt) in the presence of AMP-PNP but much weaker and more unstable in the presence of ADP (Figs S3B-D). KIF1A(1-393)(R11Q)::LZ::mSca did not stably bind to microtubules even in the presence of AMP-PNP (Figs S3D and E). These suggest that R11Q and R254Q affect microtubule binding by different mechanisms, which would account for the different behavior of these two mutants (see Discussion).

### Genetic screening in model worms identified a mutation that recovers the motility of human KIF1A

We performed a genetic screening and searched for mutants that recover the movement of *unc-104(R251Q)* model worms (Fig 4A). From about 10000 haploid genomes, we recovered two independent suppressors. Interestingly, genomic sequencing revealed that these two independent suppressor lines have the same mutation, UNC-104(D177N). While *unc-104(R251Q)* worms did not move well, *unc-104(D177N,R251Q)* showed much better performance in the swimming assay (Fig 4B). *unc-104(D177N, R251Q)* had clear synaptic puncta in the dorsal synaptic region, while a very small number of dorsal synaptic puncta were observed in *unc-104(R251Q)* worms (Fig S4A-D). UNC-104(D177) in *C. elegans* is equivalent to KIF1A(D180) in human. Then, we compared the activity of human KIF1A(R254Q) and KIF1A(D180N,R254Q) in the single molecule assays (Fig 4C-F). We found three motility parameters changed in KIF1A(R254Q) were recovered by the additional KIF1A(D180N) mutation (Fig 4E-G). The data suggest phenotypes in worm model of KAND is relevant to the activity of human KIF1A motor.

**Figure 4.**
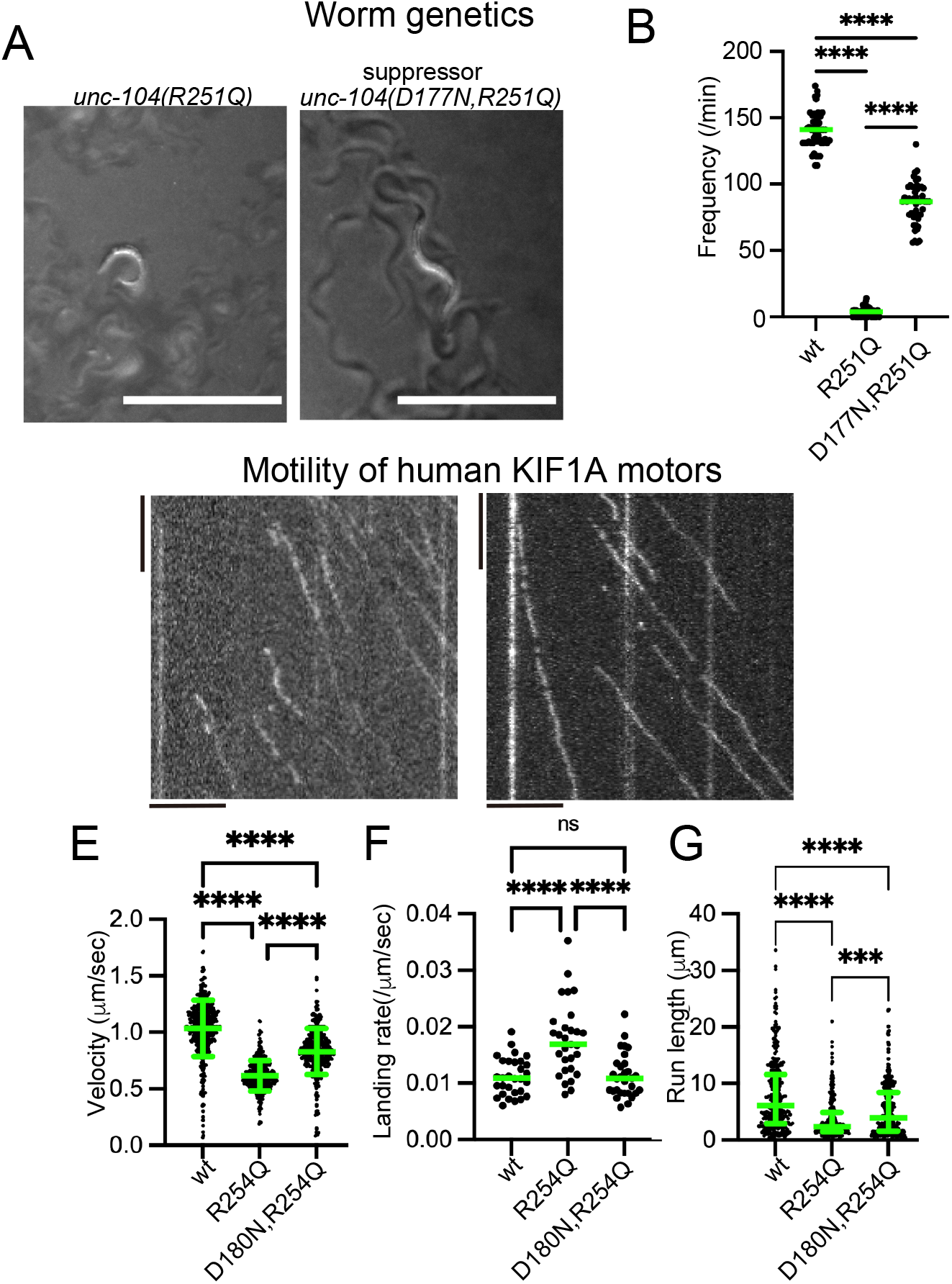
Suppressor screening. (A) Macroscopic phenotypes of *unc-104(R251Q)* and a suppressor mutant *unc-104(D177N, R251Q)*. While *unc-104(R251Q)* worms do not move well on the bacterial feeder, *unc-104(D177N, R251Q)* worms move smoothly. Bars, 1 mm. (B) Dot plots showing the result of swim test. The number of body bent in a water drop was counted for 1 min and plotted. Dots represents the number of bends from each worm. Green bars represent median value. Kruskal-Wallis test followed by Dunn’s multiple comparison test. N = 20 worms for each genotype. ****, Adjusted P value < 0.0001. (C and D) Representative kymographs showing the motility of 10 pM human KIF1A(R254Q) protein (C) and KIF1A(D180N, R254Q) protein (D) on microtubules. Vertical and horizontal bars represent 5 sec and 5 *μ*m, respectively. (E) Dot plots showing the velocity of KIF1A. Dot shows actual value from each data point. Green bars represent mean ± S.D.. Kruskal-Wallis test followed by Dunn’s multiple comparison test. n = 433 (wt), 325 (R254Q) and 368 (D180N, R254Q). ****, Adjusted P Value < 0.0001. (F) Dot plots showing the landing rate of KIF1A. The number of KIF1A that bound to microtubules was counted and adjusted by the time window and microtubule length. Green bars represent median value. Kruskal-Wallis test followed by Dunn’s multiple comparison test. n = 30 (10 pM wt), 29 (10 pM R254Q), 30 (10 pM D180N, R254Q). ****, Adjusted P Value < 0.0001. (G) Dot plots showing the run length of KIF1A. Green bars represent median value and interquartile range. Kruskal-Wallis test followed by Dunn’s multiple comparison test. n = 312 (wt), 241 (R254Q) and 312 (D180N, R254Q) homodimers. ****, Adjusted P Value < 0.0001. Note that the reported run lengths are an underestimation of the motor’s processivity. as described in Figure 3J and that KIF1A(wt) and KIF1A(R254Q) values are the same with Figure 3. See also supplementary Figure S4.

### Synaptic vesicles are mislocalized in heterozygous worms

KAND mutations, including KIF1A(R11Q), KIF1A(R254Q) and KIF1A(P305L) studied here, are *de novo* and cause neuropathies in an autosomal dominant manner. Moreover, KAND is a progressive disease. We therefore analyzed neuronal phenotypes of heterozygous worms in late adult stages (Fig 5A-F). Reverse transcription-polymerase chain reaction (RT-PCR) followed by restriction enzyme digestion confirmed that expression level of wild type and mutant *unc-104* mRNA was almost 1:1 (Fig S1A and S5A). We included a completely null allele of unc-104, *unc-104(tm819)*, as a control. *unc-104(tm819)* have a deletion mutation in the motor-domain coding region. *unc-104(tm819)* homozygotes were lethal but *unc-104(tm819)/+* was viable. *unc-104(tm819)* is described as *unc-104(null)* in this paper. DA9 synapses and body movement in the water were analyzed in heterozygotes at 1 day, 3 days, 6 days and 9 days after the final molt (Fig 5A-F and S5B-I). No significant differences were observed at 1-day adults (Figs S5B-D). At day 3, some disease-associated heterozygotes showed mislocalization of synaptic vesicles in the dendrite (Figs 5A, B and S5E-G). At day 6 and 9, dendritic mislocalization was clearly observed in 45 to 70 % *unc-104(R9Q)/+, unc-104(R251Q)/+* and u*nc-104(P298L)/+* (Fig 5C-E). In contrast, the mislocalization of synaptic puncta in *unc-104(null)/+* was comparable to *wild type* controls in all age adults (Fig 5E, S5C and S5F). More than 70% wild type and *unc-104(null)/+* worms did not show misaccumulation of GFP::RAB-3 in the dendrite even at 6 and 9 days (Fig 5E). The number of dorsal axonal puncta were not significantly affected in all age adults in all genotypes (Figs S5 B, E, H and I). Similar to the dendritic mislocalization phenotypes, worm movement was slightly affected in 6-days or 9-days adults in *unc-104(R9Q)/+, unc-104(R251Q)/+* and u*nc-104(P298L)/+* (Fig 5F). The defect is more evident at day 9. In contrast, the body movement of unc-*104(null)/+* was comparable to age-matched wild type worms (Fig 5F). Overall, phenotypes of *unc-104(R9Q)/+, unc-104(R251Q)/+* and *unc-104(P298L)/+* were heavier than *unc-104(null)/+*.

**Figure 5.**
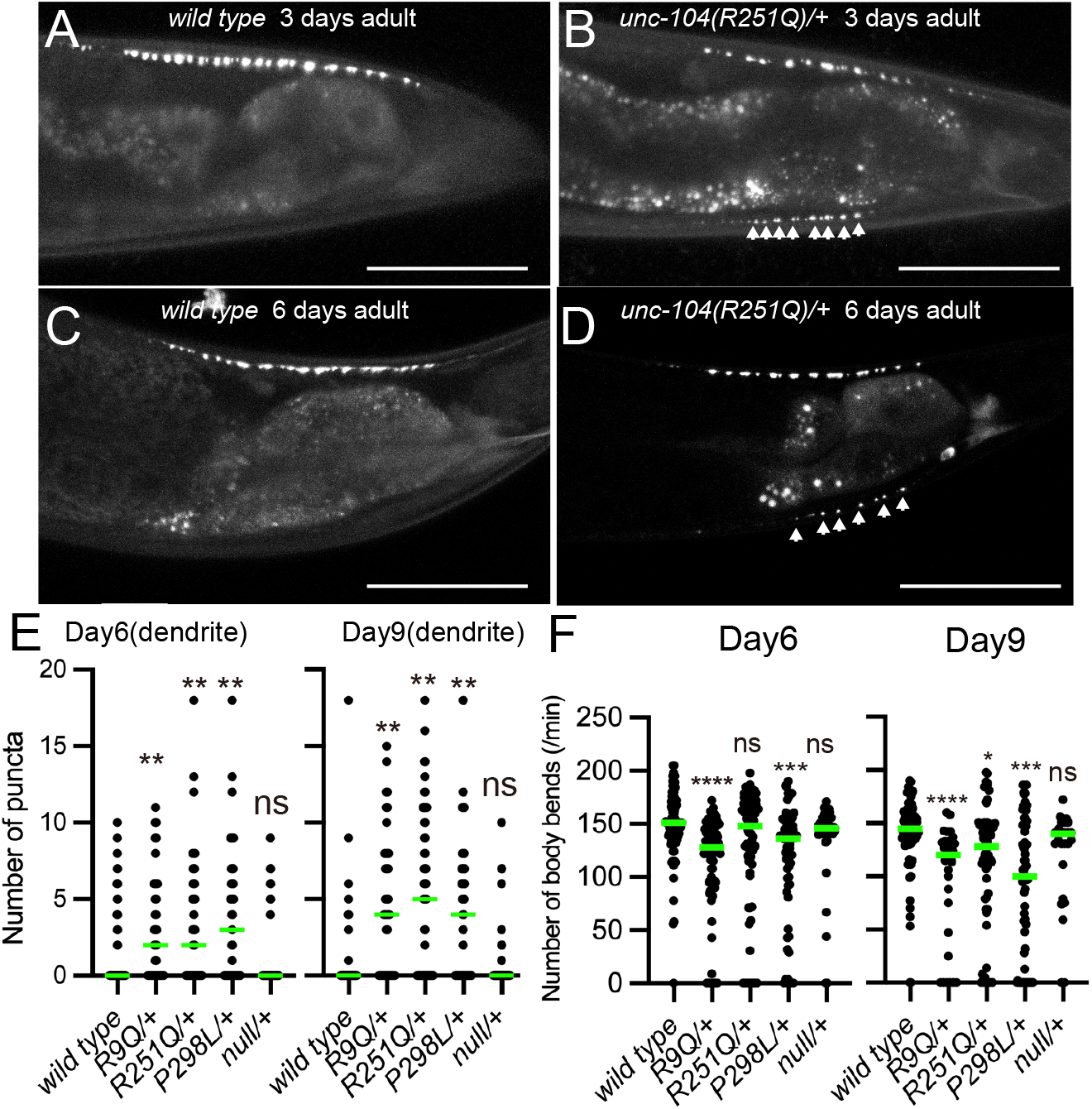
Synaptic vesicle localization of heterozygotes. (A-D) Representative images showing synaptic vesicle distribution in 3 days *wild-type* adult (A), 3 days *unc-104(R251Q)/+* adult (B), 6 days *wild-type* adult (C), and 6 days *unc-104(R251Q)/+* adult (D). Synaptic vesicles are visualized by GFP::RAB-3. Arrow heads show mislocalization of synaptic vesicles in the dendrite. Bars, 50 *μ*m. (E) Dot plots showing the number of dendritic puncta at 6 and 9 days. Each dot shows the number of puncta in the dendrite in each worm. Green bars represent median value. Kruskal-Wallis test followed by Dunn’s multiple comparison test. N = 59 (wt), 36 (R9Q/+), 46 (R251Q/+), 39 (P298L/+) and 40 (null/+) (6-days adult worms); 58 (wt), 38 (R9Q/+), 49 (R251Q/+), 40 (P298L/+) and 43 (null/+) (9-days adult worms). ns, Adjusted P Value > 0.05 and statistically not significant. **, Adjusted P Value < 0.01. (F) Dot plots showing the result of swim test at 6 and 9 days. Each dot shows the number of bends in each measurement. Green bars represent median value. Kruskal-Wallis test followed by Dunn’s multiple comparison test. N = 76 (wt), 87 (R9Q/+), 74 (R251Q/+), 65 (P298L/+) and 38 (null/+) (6-days adult worms); 66 (wt), 30 (R9Q/+), 65 (R251Q/+), 67 (P298L/+) and 27 (null/+) (9-days adult worms). ns, Adjusted P Value > 0.05 and statistically not significant. *, Adjusted P Value < 0.05. **, Adjusted P Value < 0.01. ****, Adjusted P Value < 0.0001. See also supplementary Figure S5.

### Reduced Axonal transport in disease-associated heterozygote worms

DA9 axon and dendrite have plus-end out and minus-end out microtubules, respectively (41). We analyzed axonal and dendritic transport at Day 1 to exclude a possibility that disrupted neuronal morphology indirectly reduce transport parameters. In *wild type, unc-104(R9Q)/+, unc-104(R251Q)/+* and *unc-104(P298L)/+* and *unc-104(null)/+* worms, both anterograde and retrograde movement of synaptic vesicle precursors was observed in the DA9 axon (Fig 6A-E). Vesicular movement in heterozygous worms was much better than that in homozygous worms (Fig 2). However, in three disease-associated mutant heterozygotes, the velocity of anterograde axonal transport was reduced (Fig 6B). No significant difference in anterograde velocity was detected in *unc-104(null)/+* worms. In contrast, retrograde velocity, which depends on dynein motor, was not significantly changed in all mutant heterozygotes (Fig 6C). The frequency of both anterograde and retrograde axonal transport was reduced in disease-associated mutant heterozygotes compared with than in wild type (Fig 6D and E). The directionality of vesicular transport was not significantly changed in disease-associated mutants (Fig 6F). These results are consistent with previous studies showing that inhibition of anterograde machineries affect retrograde transport machineries, and vice versa (37, 42-44). In the dendrite, even though misaccumulation of stable puncta was not observed in *wild type*, some motile vesicles could be detected as described (37) (Fig S6). While the velocity of retrograde transport (i.e. transport from the dendritic tip to the cell body) was slightly reduced (Fig S6B), other parameters were not strongly affected (Fig. S6). This would be because multiple motors transport synaptic vesicle precursors in dendrite (37). These data are consistent with the previous mathematical model showing that misaccumulation of synaptic vesicles to dendrite is caused mainly by reduced anterograde transport in the proximal axon (37).

**Figure 6.**
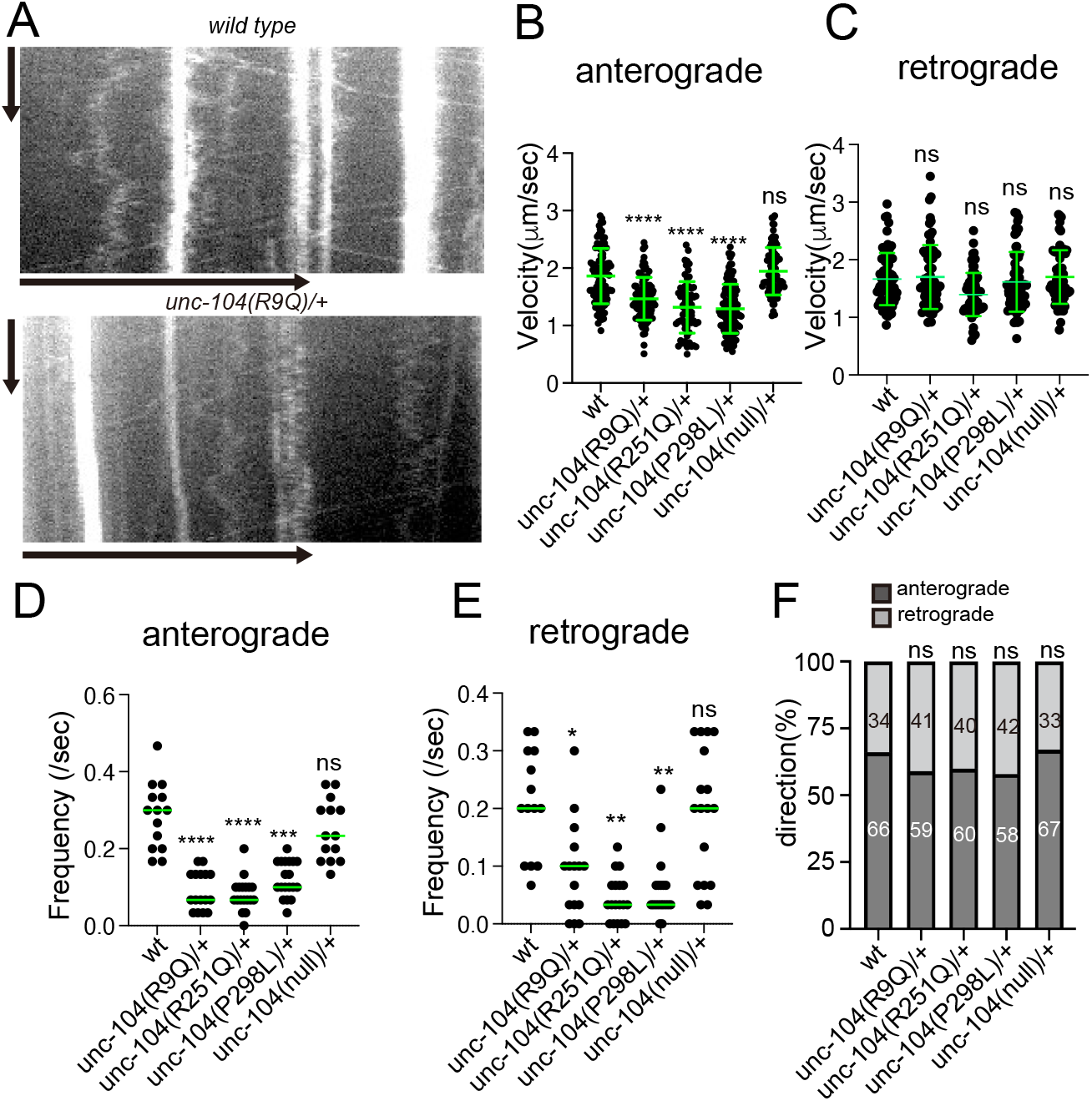
Axonal transport in KAND model heterozygotes. (A)Representative kymographs showing axonal transport of synaptic vesicle precursors in *wild type* and *unc-104(R9Q)/+* at 1 day adults. GFP::RAB-3 was used as a marker. Vertical and Horizontal bars show 10 seconds and 10 *μ*m, respectively. (B and C) The velocity of axonal transport. The velocity of anterograde transport (B) and retrograde transport (C) are shown as dot plots. (B) Kruskal-Wallis test followed by Dunn’s multiple comparison test. Green bars show mean ± S.D.. n = 94 (wild type), 90 (R9Q/+), 66 (R251Q/+), 117 (P298L/+) and 84 (null/+) vesicles from at least 5 independent worms. ns, Adjusted P Value > 0.05 and no significant statistical difference. ****, Adjusted P Value < 0.0001. (C) Kruskal-Wallis test followed by Dunn’s multiple comparison test. Green bars show mean ± S.D.. n = 70 (wild type), 70 (R9Q/+), 68 (R251Q/+), 63 (P298L/+) and 65 (null/+) vesicles from at least 5 independent worms. ns, Adjusted P Value > 0.05 and no significant statistical difference. (D and E) Frequency of axonal transport. The frequency of anterograde transport (D) and retrograde transport (E) are shown as dot plots. (E) Kruskal-Wallis test followed by Dunn’s multiple comparison test. Each dot represent data from each worm. Green bars represent median value. N = 14 (wt), 16 (R9Q/+), 18 (R251Q/+) and 19 (P298L/+) independent worms. ****, Adjusted P Value < 0.0001. (E) Kruskal-Wallis test followed by Dunn’s multiple comparison test. Each dot represent data from each worm. Green bars represent median value. N = 14 (wt), 16 (R9Q/+), 18 (R251Q/+) and 19 (P298L/+) independent worms. **, Adjusted P Value < 0.01, ****, Adjusted P Value < 0.0001. (F) Directionality of vesicle movement. The number in the bar graph shows the actual percentage. ns, Adjusted P Value > 0.05 and statistically not significant. Chi-square test. Compared to wt worms. See also supplementary Figure S6.

### Diesease mutant/wild type heterodimers have reduced motor properties

The KIF1A motor forms a homodimer for efficient anterograde axonal transport (34). In patients who have autosomal dominant mutations, half of the motor complex in the neuron is expected to be heterodimers composed of wild-type KIF1A and disease-associated KIF1A. But the behavior of heterodimers on microtubules remains largely unanalyzed. To analyze the motility of heterodimers at a single-molecule resolution, we purified heterodimers composed of wild-type KIF1A and disease-associated KIF1A. Wild-type KIF1A fused with leucine zipper and mScarlet-I (KIF1A(1-393)::LZ::mSca) and disease-associated KIF1A without fluorescent tag (KIF1A(1-393)::LZ) were co-expressed in bacteria (Fig 7A). The two constructs were respectively fused with Strep tag and His tag for purification. Tandem affinity purification using His tag and Strep tag followed by gel filtration was performed to purify heterodimers. We analyzed heterodimers composed of KIF1A(1-393)::LZ::mSca and KIF1A(1-393)::LZ that were recovered from the same peak fractions (Fig 7B and supplementary Fig S7). The ratio between two subunits calculated from band intensities and molecular weight were about 1:1, indicating heterodimers.

**Figure 7.**
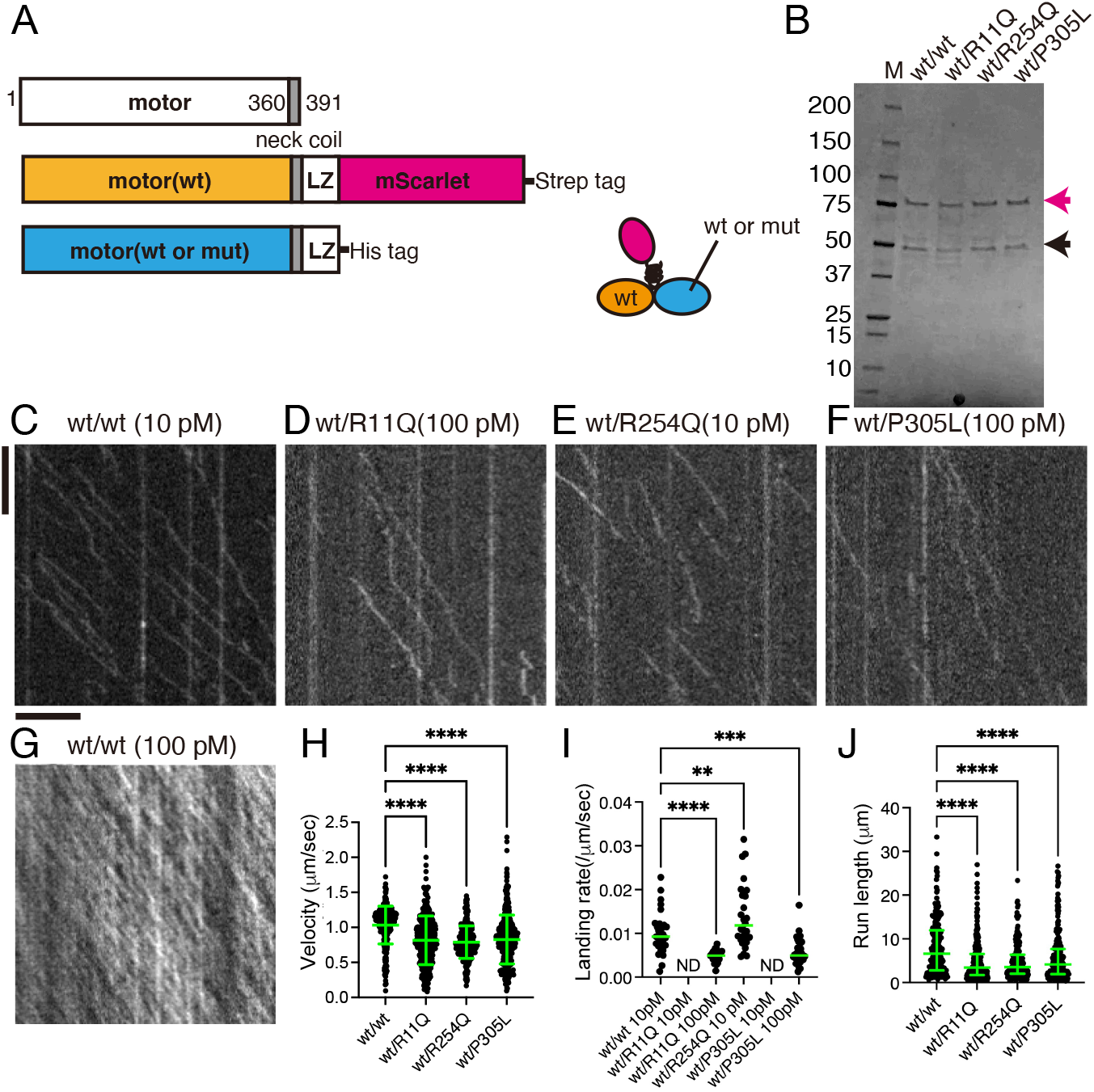
The single molecule behavior of wild type/mutant KIF1A heterodimers. (A) Schematic drawing of the recombinant KIF1A heterodimer analyzed in Figure 7. (B) Purified KIF1A(1–393)::LZ::mScarlet/KIF1A(1–393)::LZ heterodimers were separated by SDS-PAGE and detected by Coomassie brilliant blue staining. M represents marker. Numbers on the left indicate the molecular weight (kDa). Magenta and black arrows indicate KIF1A(1–393)::LZ::mScarlet and KIF1A(1–393)::LZ, respectively. (C-G) Representative kymographs showing the motility of 10 pM KIF1A (wt) (C), 100 pM KIF1A(R11Q) (D), 10 pM KIF1A(R254Q) (E), 100 pM KIF1A(P305L) and 100 pM KIF1A (wt) (G). Vertical and horizontal bars represent 5 sec and 5 *μ*m, respectively. (H) Dot plots showing the velocity of KIF1A. Each dot shows each data. Green bars represent mean ± S.D.. Kruskal-Wallis test followed by Dunn’s multiple comparison test. n = 308 (wt/wt), 315 (wt/R11Q), 294 (wt/R254Q) and 414 (wt/P305L) heterodimers. ****, Adjusted P Value < 0.0001. (I) Dot plots showing the landing rate of KIF1A. The number of KIF1A that binds to microtubules was counted and adjusted by the time window and microtubule length. Each dot shows each data. Green bars represent median value. Kruskal-Wallis test followed by Dunn’s multiple comparison test. n = 29 (10 pM wt/wt), 29 (100 pM wt/R11Q), 28 (10 pM wt/R254Q) and 38 (100 pM wt/P305L) independent observations. **, Adjusted P Value < 0.01, ***, Adjusted P Value < 0.001, ****, Adjusted P Value < 0.0001. (J) Dot plots showing the run length of KIF1A. Each dot shows each data. Green bars represent median value and interquartile range. Kruskal-Wallis test followed by Dunn’s multiple comparison test. n = 215 (wt/wt), 241 (wt/R11Q), 195 (wt/R254Q) and 266 (wt/P305L) heterodimers. ****, Adjusted P Value < 0.0001. Note that the reported run lengths are an underestimation of the motor’s processivity as described in Figure 3J. See also supplementary Figure S7.

As a positive control, we compared the motility of KIF1A(1-393)::LZ::mSca/KIF1A(1-393)::LZ heterodimers with KIF1A(1-393)::LZ::mSca homodimers (Fig 3C and 7C). Velocity, landing rate and run length of wild-type homodimers and heterodimers were statistically the same (Velocity: 1.03 ± 0.24 µm/sec and 1.03 ± 0.26 µm /sec, Run length: 7.99 ± 6.42 µm and 8.07 ± 6.30 µm, Landing rate: 0.011 ± 0.003 µm ^-1^s^-1^ and 0.010 ± 0.004 µm ^-1^s^-1^ for homodimers and heterodimers, respectively. Mean ± standard deviation. Statistically not significant by t-test.). In contrast, heterodimers composed of wild-type KIF1A and disease-associated KIF1A showed reduced motility (Fig 7C-J). Although Although KIF1A(1–393)(R11Q)::LZ::mSca showed no processive movement on microtubules, KIF1A(1–393)::LZ::mSca/KIF1A(1–393)(R11Q)::LZ heterodimers showed processive movement (Fig 7D). Other two heterodimers also showed processive movement (Figs 7E and F). Then, we analyzed physical parameters of these heterodimers in detail. The velocity of KIF1A(1–393)::LZ::mSca/KIF1A(1–393)(R11Q)::LZ, KIF1A(1–393)::LZ::mSca/KIF1A(1–393)(R254Q)::LZ and KIF1A(1–393)::LZ::mSca/KIF1A(1–393)(P305L)::LZ heterodimers was lower than that of wild-type KIF1A (Fig 7H). The landing event of KIF1A(1–393)::LZ::mSca/KIF1A(1–393)(R11Q)::LZ and KIF1A(1–393)::LZ::mSca/KIF1A(1–393)(P305L)::LZ heterodimers on microtubules could not be observed at 10 pM (Fig 7I). At 100 pM, in which wild-type KIF1A homodimers were saturated on microtubules (Fig 7G), the motility of KIF1A(1–393)::LZ::mSca/KIF1A(1–393)(R11Q)::LZ and KIF1A(1–393)::LZ::mSca/KIF1A(1–393)(P305L)::LZ dimers was observed (Fig 7D, F and I) but the run lengths of these wild-type/mutant dimers were much shorter compared with that of wild-type dimers (Fig 7J). The landing rate of KIF1A(1–393)::LZ::mSca/KIF1A(1–393)(R254Q)::LZ heterodimers was higher than that of wild-type dimers (Fig 7I). However, run length of KIF1A(1–393)LZ-mSca/KIF1A(1–393)(R254Q)LZ heterodimers was shorter than that of wild-type dimers (Fig 7J). These results show that KAND mutations strongly affect the landing rate and motility parameters in heterodimers with wild-type KIF1A.

### Dominant negative effects of disease-associated mutations *in vitro* and *in vivo*

Multiple kinesin dimers cooperatively transport cargo vesicles in the cell (30, 45, 46, 47). Thus, it is expected that the ratio of wt/wt homodimers, wt/mutant heterodimers and mutant/mutant homodimers is 1:2:1 on cargo vesicles in KAND patients who have heterozygous mutations. To mimic the condition, we performed microtubule gliding assays using mixed motors (45, 48) (Fig 8A). The velocity of wt/mutant heterodimers and mutant/mutant homodimers were significantly reduced in the gliding assay (Fig 8B), that is consistent with the results of single molecule assays. In the mixed condition, all three mutants inhibited the motility of wild type KIF1A (Fig 8C). As shown previously, reduced concentration of KIF1A(wt) protein does not significantly affect the velocity in the gliding assay (Fig S8A) (45, 48, 49). Thus, the reduced velocity observed in the mixed condition is thought to be an inhibitory effect of mutant motors. The microtubule gliding velocity showed 3-40% reduction, which is similar to the slower anterograde transport in heterozygous worms (Fig 6B). Finally, to show that KAND mutations dominant negatively inhibit the axonal transport *in vivo, unc-104(R9Q), unc-104(R251Q) and unc-104(P298L)* cDNA, corresponding to KIF1A(R11Q), KIF1A(R254Q) and KIF1A(P305L) mutants, were overexpressed in DA9 neuron (Fig 8D-G and S8B and S8C). As a result, in 70% UNC-104(R9Q), UNC-104(R251Q) and UNC-104(P298L)-expressed animals, synaptic vesicles were misaccumulated to the proximal region of the DA9 axon (Fig 8B, C and S8). No significant effects were observed in UNC-104(wt)-overexpressed worms. These *in vitro* and *in vivo* data suggests that all three mutations analyzed in this study reduces axonal transport by dominant-negative manner.

**Figure 8.**
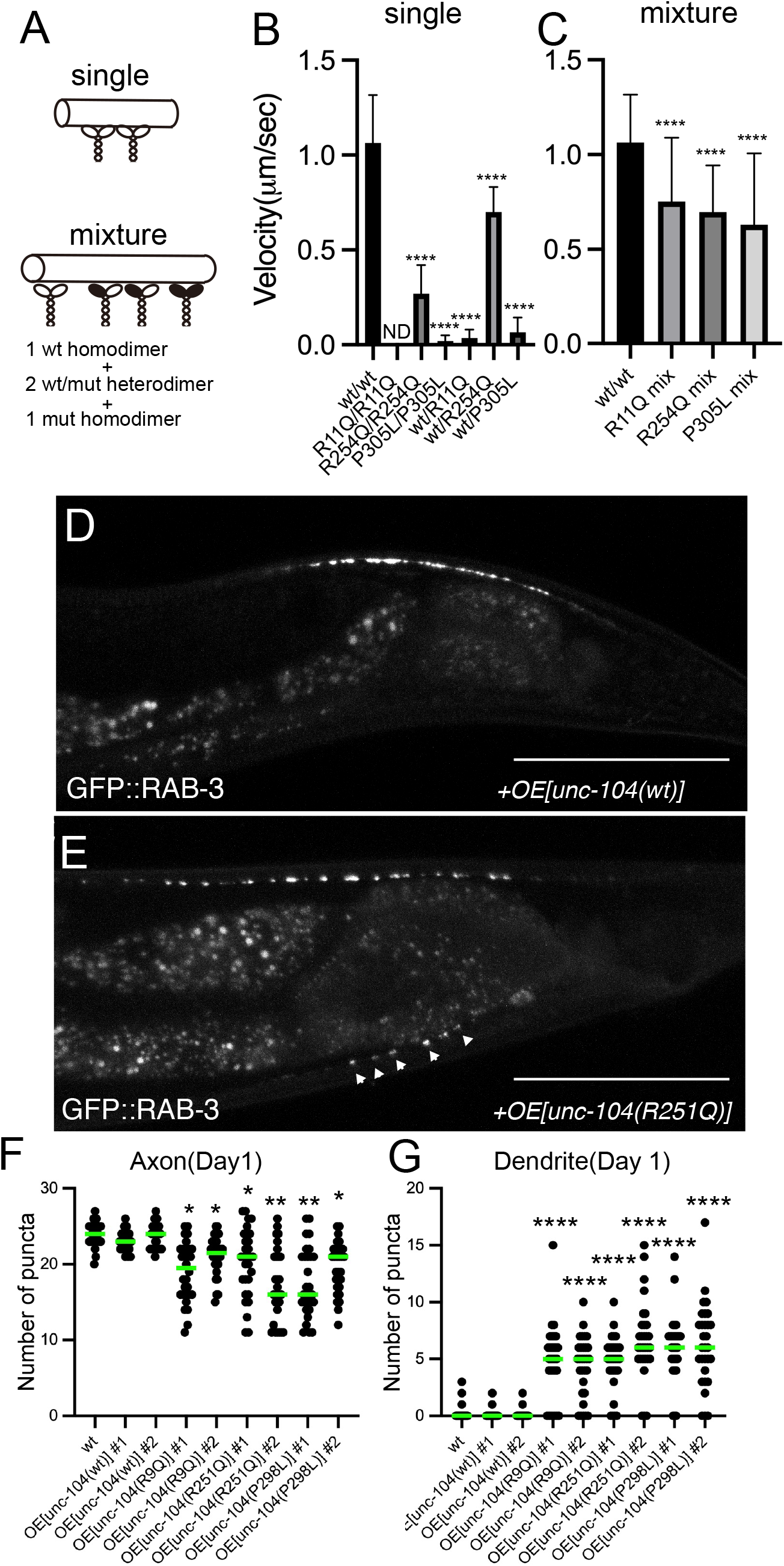
Dominant negative nature of KAND mutations. (A-C) Microtubule gliding assays. Shematic drawing of the microtubule gliding assay in different conditions (A). (B) Microtubule gliding assays using single motors. Bars and error bars represent mean and SD, respectively. N = 146 (30 nM wt homodimers), 112 (30 nM R254Q homodimers), 20 (100 nM P305L homodimers), 59 (30 nM wt/R11Q heterodimers), 130 (30 nM wt/R254Q heterodimers) and 43 (30 nM wt/P305L heterodimers) microtubules from at least three independent experiments. Note that no microtubule movement was observed in 100 nM KIF1A(R11Q) homodimers. Kruskal-Wallis test followed by Dunn’s multiple comparison test.****, p < 0.0001. (C) Microtubule gliding assays using mixed motors. Bars and error bars represent mean and SD, respectively. N = 146 (30 nM wt), 102 (30 nM R11Q mixture), 134 (30 nM R254Q mixture) and 108 (30 nM P305L mixture) microtubules. Kruskal-Wallis test followed by Dunn’s multiple comparison test. ****, p < 0.0001. (D-G) UNC-104(wt), UNC-104(R9Q), UNC-104(R251Q) and UNC-104(P298L) were overexpressed in the wild-type background and the localization of synaptic vesicles was observed. (D and E) Representative images showing the localization of synaptic vesicles in UNC-104(wt)-expressing worm (D) and UNC-104(R251Q)-expressing worm (E). Arrowheads show synaptic-vesicle accumulated puncta that are mislocalized in the dendrite. Bars, 50 *μ*m. (F and G) Dot plots showing the number of ventral axonal and dorsal dendritic puncta at 1 day. Each dot shows the number of puncta in the dorsal axon (F) and ventral dendrite (G) in each worm. Green bars represent median value. N = 30 worms from each strain. Kruskal-Wallis test followed by Dunn’s multiple comparison test. ns, Adjusted P Value > 0.05 and no significant statistical difference. *, Adjusted P Value < 0.05. **, Adjusted P Value < 0.01. ****, Adjusted P Value < 0.0001. See also supplementary Figure S8.

## Discussion

It is important to determine heterozygous disease-associated mutations are dominant negative or haploinsufficiency because the difference significantly affects treatment strategies. Our data suggest *de novo* and autosomal dominant KAND mutations perturb axonal transport by two mechanisms. One inhibitory mechanism is induced by heterodimerization. Axonal transport motors form homodimers that move processively on microtubules (50). When a mutation in a motor protein gene is dominant, and if the mutation does not affect the stability, expression or activation of the motor protein, half of the motor dimers in the cell are predicted to be heterodimers composed of wild-type motor and disease-associated motor. Many disease-associated mutations in motor proteins are caused by autosomal dominant mutations; however, little attention has been paid to the properties of heterodimers in motor-associated diseases and previous studies have mainly analyzed the properties of mutant homodimers *in vitro* (26, 27, 51, 52). We show here disease-associated KIF1A inhibits the motility of wild type KIF1A by forming dimers. Another inhibitory effect caused by disease-associated KIF1A is caused when KIF1A motors work as a team. In the axon, multiple motor dimers bind to and cooperatively transport a vesicle (Fig 9) (46, 47). Microtubule gliding assays performed using mixed motors show that cooperative transport is inhibited when disease-associated heterodimers and homodimers are mixed. Overexpression of mutant *unc-104* cDNAs dominant-negatively induces mislocalization of synaptic vesicles in wild-type neuron (Figs 8D-G). Together with the data showing that *unc-104(R9Q)/+, unc-104(R251Q)/+* and *unc-104(P298L)/+* but not *unc-104(null)/+*, show defects in axonal transport, we suggest that these disease-associated mutations cause neuronal symptoms mainly by dominant-negative mechanisms (Fig 9). *In vitro* assays show KIF1A(R11Q) motor does not move on microtubules at all but KIF1A(R254Q) and KIF1A(P305L) motors move to some extent (Fig 3 and Fig 8B). Similar huge variation has been observed in the activity of mutant homodimers with other mutations (27). However, degree of severity in some KAND patients, as well as model worms, are not always consistent with properties of mutant homodimers (27-29). These may be explained by the dominant negative nature of mutations. This is partially supported by the microtubule gliding assay in which the difference in the mixed condition is much smaller than that of in single mutant homodimers (Fig 8). Mutations in other axonal transport motors, such as KIF5A and cytoplasmic dynein heavy chain 1 genes, are causes of autosomal dominant neuropathies (51-53). Similar phenomena observed here may underly in the pathogenesis of these neuropathies.

**Figure 9.**
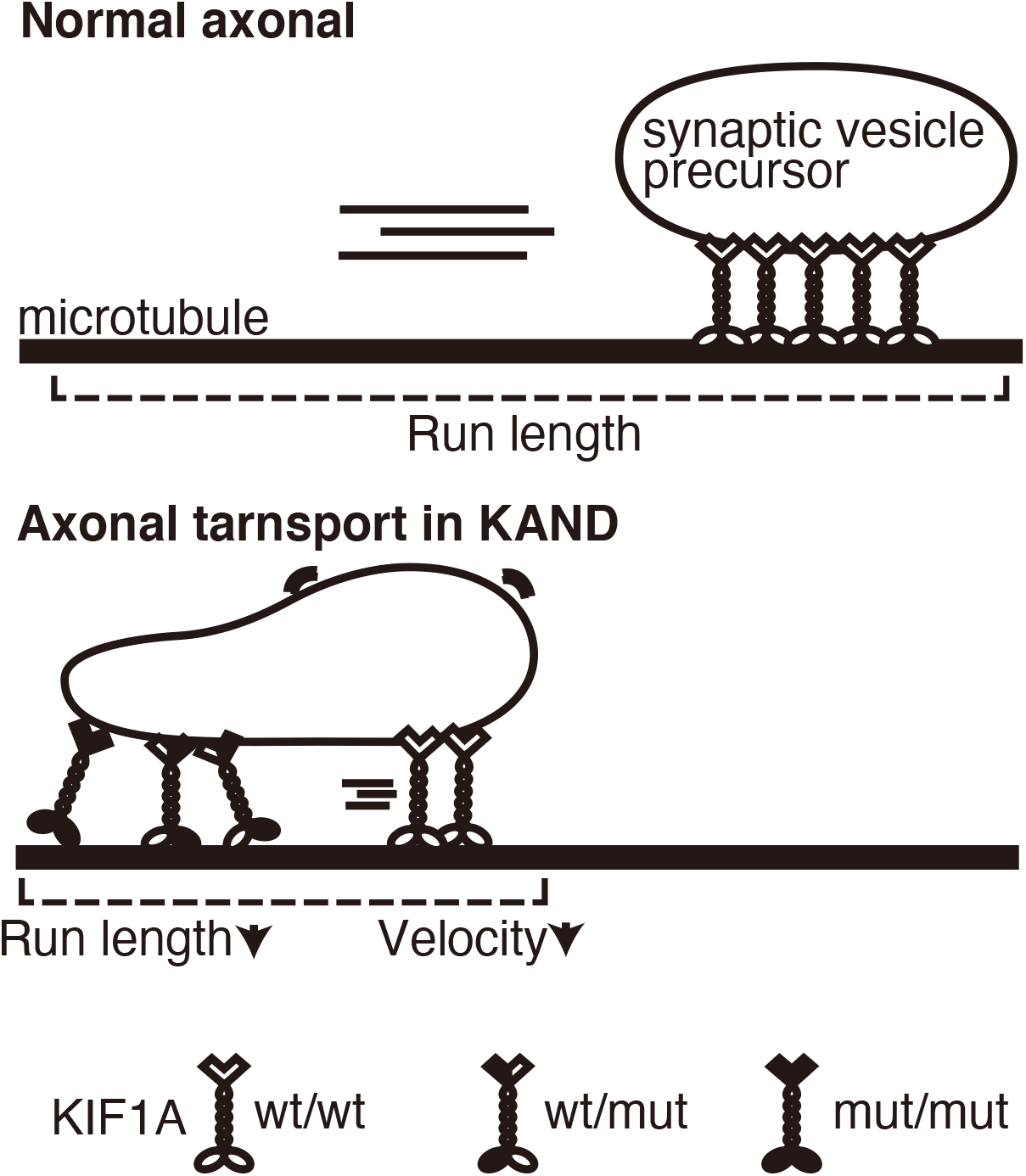
Model. Schematic drawing showing how vesicular transport is suppressed in KAND patient axons. Not only mutant homodimers but also wild-type/mutant heterodimers inhibit axonal transport of synaptic vesicle precursors.

We tested three mutations, R11Q, R254Q and P305L. It has been shown that KIF1A(P305) is in the L12 loop which supports the binding with microtubules (32). Consistent with this, we detected reduced microtubule binding in KIF1A(P305L) (Fig 3). We show here that KIF1A(R11Q) reduces the binding frequency with microtubules while KIF1A(R254Q) reduces run length (Fig 3). This difference would be caused because these two mutations affect different nucleotide states of KIF1A. The affinity of KIF1A to the microtubules undergoes a drastic change during the hydrolysis of ATP into ADP (12). KIF1A alternates between a strong binding state (ATP binding state) and a weak binding state (ADP binding state) with microtubules. R11 is at the terminal of *β*1 sheet that stabilizes ATP binding pocket, suggesting that the R11Q mutation would affect the ADP release or binding with new ATP. Another possibility is the R11Q mutation destabilizes the folding of the motor domain. R254 is in the L11 loop. The L11 loop stabilizes *α*4 helix, a positively-charged microtubule-binding interface, in the weak binding state (ADP state) (54). Thus, R254Q mutation would destabilize the *α*4 helix in the weak binding state (ADP state) and cause more frequent dissociation from microtubules during ATP hydrolysis, leading to the short run length. Consistent with these ideas, the microtubule binding of KIF1A(R11Q) is very weak even in the presence of AMP-PNP (Fig S6). In contrast, the microtubule binding of KIF1A(R254Q) is weaker than KIF1A(wt) in the presence of ADP but comparable to KIF1A(wt) in the presence of AMP-PNP (Fig S6). Because the suppressor mutation (D180N) reduces the negative charge on the motor surface, the mutation would strengthen the binding of KIF1A(R254Q) with microtubules and would recover the run length (Fig 4).

Multiple kinesin dimers cooperatively transport cargo vesicles. Force generation is fundamental to the cooperative transport (30, 45, 46). Thus, one concern is whether the single molecule assays performed in unloaded conditions is relevant to the axonal transport of synaptic vesicle precursors. Unbiased genetic screening using the model worm identified the *unc-104(D177N)* mutation recovers the axonal transport in *unc-104(R251Q)* worms (Fig 4A and B and S4). An orthologous mutation, KIF1A(D180N), significantly recovered the motility of human KIF1A(R254Q) in the single molecule assays (Fig 4C-G). These suggest the activity measured by the single molecule assay is relevant to the axonal transport *in vivo*. The data also suggest worm models established here reflect the transport activity of human KIF1A. As far as we searched, almost all motor domain residues that are mutated in KAND are conserved in *C. elegans* UNC-104. Thus, worm models, including three lines established in this work, would be the foundation for future genetic screenings and drug screenings to search for treatment of KAND.

## Methods

### Worm experiments

*C. elegans* strains were maintained as described previously (55). N2 wild-type worms and OP50 feeder bacteria were obtained from the *C. elegans* genetic center (CGC) (Minneapolis, MN, USA). Transformation of *C. elegans* was performed by DNA injection as described (56). The swim test was performed as described previously (57).

### Genome editing

Target sequences for cas9 and repair templates used to make *unc-104* mutants are described in ***supplementary table S1***. Target sequences were inserted into pRB1017 (a gift from Andrew Fire, Stanford University, addgene #59936). pDD162 (a gift from Bob Goldstein, UNC Chapel Hill, addgene #47549) was used to express Cas9. These vectors and oligonucleotides were injected into young adult worms as described with a slight modification (35). Genotype was confirmed by PCR followed by Sanger sequencing.

### Strains

Strains used in this study are described in ***supplementary table S2***. Heterozygotes that have the *wyIs251* marker were generated by crossing *unc-104* homozygotes *wyIs251* males. Heterozygotes with wyIs85 markers were maintained by *mIn1* balancer. F1 worms showing non-unc phenotypes at the L4 stage were picked and transferred to new plates. Next day, adult worms were transferred to new plates (Day 0). Worms were transferred to new plates until the observation.

### Statistical analyses and graph preparation

Statistical analyses were performed using Graph Pad Prism version 9. Statistical methods are described in the figure legends. Graphs were prepared using Graph Pad Prism version 9, exported in the TIFF format and aligned by Adobe Illustrator 2021.

### Purification of recombinant KIF1A

Reagents were purchased from Nacarai tesque (Kyoto, Japan), unless described. Plasmids to express recombinant KIF1A are described in ***supplementary table S3***. Proteins were expressed in BL21(DE3) and purified by Streptactin-XT resin (IBA Lifesciences, Göttingen, Germany) in the case of homodimers and Streptactin-XT resin and TALON resin (Takara Bio Inc., Kusatsu, Japan) in the case of heterodimers. Eluted fractions were further separated by NGC chromatography system (Bio-Rad) equipped with a Superdex 200 Increase 10/300 GL column (Cytiva).

### TIRF single-molecule motility assays

TIRF assays were performed as described (25). Tubulin was purified from porcine brain as described (58). Tubulin was labeled with Biotin-PEG_2_-NHS ester (Tokyo Chemical Industry, Tokyo, Japan) and AZDye647 NHS ester (Fluoroprobes, Scottsdale, AZ, USA) and polymerized as described (59). An ECLIPSE Ti2-E microscope equipped with a CFI Apochromat TIRF 100XC Oil objective lens, an Andor iXion life 897 camera and a Ti2-LAPP illumination system (Nikon, Tokyo, Japan) was used to observe single molecule motility. NIS-Elements AR software ver. 5.2 (Nikon) was used to control the system.

## Data Availability

All study data are included in the article and/or supporting information.

## Acknowledgements

YA was supported by the Advanced Graduate Program for Future Medicine and Health Care, Tohoku University. KH was supported by JST PRESTO (grant No. JPMJPR1877) and FRIS Creative Interdisciplinary Research Program, Tohoku University. SN was supported by JSPS KAKENHI (20H03247, 19H04738, 20K21378), the Naito foundation and the Uehara foundation. Some worm strains and OP50 were obtained from the CGC. *unc-104(tm819)/mIn1* was obtained from the NBRP. We thank Jeremy Allen, PhD, from Edanz (https://jp.edanz.com/ac) for editing a draft of this manuscript.

