## Supplementary Information for "*De novo* mutations in KIF1A-associated neuronal disorder (KAND) dominant-negatively inhibit motor activity and axonal transport of synaptic vesicle precursors"

***Supplementary text***

***Methods***

***Worm experiments***

*C. elegans* strains were maintained as described previously (1). Wild-type worm N2 and feeder bacteria OP50 were obtained from *C. elegans* genetic center (CGC) (Minneapolis, MN, USA). Nematode Growth Media (NGM) agar plates were prepared as described (1). Transformation of *C. elegans* was performed by DNA injection as described (2).

***Genome editing***

Target sequences for cas9 and repair templates used to make *unc-104* mutants are described in Figure S1A and ***supplementary table S1***. Target sequences were inserted to pRB1017 (a gift from Andrew Fire, Stanford University, addgene #59936). pDD162 (a gift from Bob Goldstein, UNC Chapel Hill, addgene #47549) was used to express Cas9. These vectors and oligonucleotides were injected to young adult worms as described with a slight modification (3). For *unc-104(R251Q)*, 50 ng of PDD162, 50 ng of unc-104(A252V)#4 and 0.6 µM of repair template were mixed and injected. Strong unc worms in the next generation was directly picked and genotyped by PCR. For *unc-104(R9Q)* and *unc-104(P298L)*, 50 ng of pDD162, 50 ng of sgRNA expression plasmid for *unc-104*, 50 ng of sgRNA expression plasmid for *ben-1*, 0.6 µM of repair template for *unc-104* were mixed and injected. Injected worms were put on NGM plate with OP50 feeder supplemented with benzoimidazole. Next generation was scored, and benzoimidazole-resistant worms were picked. These plates were genotyped by PCR followed by restriction enzyme digestion.

***Strains***

Strains used in this study is described in ***supplementary table S2***.

***Suppressor screening and determination of the suppressor mutation***

EMS mutagenesis was performed as described (4) with slight modifications. *unc-104(R251Q)*; *wyIs85* worms were treated with ethyl methanesulfonate (EMS). After the treatment, worms were washed and separated to 10 NGM plates with OP50 feeder. Plates were examined under a Zeiss dissecting microscope for improved movement in a non-clonal screen of approximately 10,000 haploid genomes. Suppressors that were picked up from different plates were considered independent. Two independent suppressors, *jpn44* and *jpn45* were recovered. We could not separate *jpn44* and *jpn45* mutations from *unc-104(R251Q)*. Then, we sequenced the genomic region of *unc-104* and found a D177N mutation.

***Analysis of the worm phenotypes***

Heterozygous worms were maintained by using *mIn1* balancer. Superficially wild type worms are heterozygotes while *unc-104* homozygotes are totally unc or lethal and *mIn1* homozygotes are dpy. To prepare age-matched worms, L4 heterozygous worms were picked up. Next day, adult worms were again transferred to new NGM plates with OP50 feeders (day 0). Worms were transferred to new plates every day until observation.

For steady state imaging of synaptic puncta, strains expressing *gfp::rab-3* under the *itr-1* promoter were analyzed at indicated days. An inverted Carl Zeiss Axio Observer Z1 microscope equipped with a 40x/1.3 objective and LSM 800 confocal scanning unit was used. Swim test was performed as described previously (5).

For time lapse imaging, worms at day 1 were observed on an inverted Carl Zeiss Axio Observer Z1 microscope equipped with a Plan-Apochromat 100x/1.4 objective and a Hamamatsu ORCA flash v3 sCMOS camera. To visualize axonal transport and dendritic transport, *gfp::rab-3* was expressed under the *mig-13* promoter and the *itr-1* promoter, respectively. Movies are taken at 4 frames per sec for 1 minutes and analyzed by Fiji software as described.

***Statistical analyses and graph preparation***

Statistical analyses were performed using Graph Pad Prism version 9. Statistical methods are described in the figure legends. Graphs were prepared using Graph Pad Prism version 9, exported in the TIFF format and aligned by Adobe Illustrator 2021.

***Purification of homodimers***

Reagents were purchased from Nacarai tesque (Kyoto, Japan), unless described. Plasmids to express recombinant KIF1A are described in ***supplementary table S3***. To purify KIF1A(1–393)::LZ::mScarlet-I::Strep homodimers, BL21(DE3) was transformed and selected on LB agar supplemented with kanamycin at 37°C overnight. Colonies were picked and cultured in 10 ml LB medium supplemented with kanamycin overnight. Next morning, 5 ml of the medium was transferred to 500 ml 2.5× YT (20 g/L Tryptone, 12.5 g/L Yeast Extract, 6.5 g/L NaCl) supplemented with 10 mM phosphate buffer (pH 7.4) and 50 µg/ml kanamycin in a 2 L flask and shaken at 37°C. Two flasks were routinely prepared. When $OD_{600}$ reached 0.6, flasks were cooled in ice-cold water for 30 min. Then, 23.8 mg IPTG was added to each flask. Final concentration of IPTG was 0.2 mM. Flasks were shaken at 18°C overnight.

Next day, bacteria expressing recombinant proteins were pelleted by centrifugation (3000 g, 10 min, 4°C), resuspended in PBS and centrifuged again (3000 g, 10 min, 4°C). Pellets were resuspended in protein buffer (50 mM Hepes, pH 8.0, 150 mM $KCH_3COO$, 2 mM $MgSO_4$, 1 mM EGTA, 10% glycerol) supplemented with Phenylmethylsulfonyl fluoride (PMSF). Bacteria were lysed using a French Press G-M (Glen Mills, NJ, USA) as described by the manufacturer. Lysate was obtained by centrifugation (75,000 g, 20 min, 4°C). Lysate was loaded on Streptactin-XT resin (IBA Lifesciences, Göttingen, Germany) (bead volume: 2 ml). The resin was washed with 40 ml Strep wash buffer (50 mM Hepes, pH 8.0, 450 mM $KCH_3COO$, 2 mM $MgSO_4$, 1 mM EGTA, 10% glycerol). Protein was eluted with 40 ml Strep elution buffer (50 mM Hepes, pH 8.0, 150 mM $KCH_3COO$, 2 mM $MgSO_4$, 1 mM EGTA, 10% glycerol, 300 mM biotin). Eluted solution was concentrated using an Amicon Ultra 15 (Merck) and then separated on an NGC chromatography system (Bio-Rad) equipped with a Superdex 200 Increase 10/300 GL column (Cytiva). Peak fractions were collected and concentrated using an Amicon Ultra 4 (Merck). Concentrated proteins were aliquoted and snap frozen in liquid nitrogen.

***Purification of heterodimers***

BL21(DE3) cells transformed with KIF1A(1–393)::LZ::mScarlet-I::Strep plasmid were cultured in LB supplemented with kanamycin at 37°C. Competent cells were prepared using a Mix&Go kit (Zymogen). The competent cells were further transformed with

KIF1A(1–393)::LZ::His plasmid and selected on LB agar supplemented with ampicillin and kanamycin. Colonies were picked and cultured in 10 ml LB medium supplemented with ampicillin and kanamycin overnight. Next morning, 5 ml of the medium was transferred to 500 ml 2.5× YT supplemented with carbenicillin and kanamycin in a 2 L flask and shaken at 37°C. Two flasks were routinely prepared. The procedures for protein expression in bacteria and preparation of bacterial lysate were the same as for the purification of homodimers. Lysate was loaded on Streptactin-XT resin (bead volume: 2 ml). The resin was washed with 40 ml wash buffer. Protein was eluted with 40 ml protein buffer supplemented with 300 mM biotin. Eluted solution was then loaded on TALON resin (Takara Bio Inc., Kusatsu, Japan)(bead volume: 2 ml). The resin was washed with 40 ml His-tag wash buffer (50 mM Hepes, pH 8.0, 450 mM $KCH_3COO$, 2 mM $MgSO_4$, 10 mM imidazole, 10% glycerol) and eluted with His-tag elution buffer (50 mM Hepes, pH 8.0, 450 mM $KCH_3COO$, 2 mM $MgSO_4$, 10% glycerol, 500 mM imidazole). Eluted solution was concentrated using an Amicon Ultra 15 and then separated on an NGC chromatography system (Bio-Rad) equipped with a Superdex 200 Increase 10/300 GL column (Cytiva). Peak fractions were collected and concentrated using an Amicon Ultra 4. Concentrated proteins were aliquoted and snap frozen in liquid nitrogen.

***TIRF single-molecule motility assays***

TIRF assays were performed as described (6). Tubulin was purified from porcine brain as described (7). Tubulin was labeled with Biotin-$PEG_2$-NHS ester (Tokyo Chemical Industry, Tokyo, Japan) and AZDye647 NHS ester (Fluoroprobes, Scottsdale, AZ, USA) as described (8). To polymerize Taxol-stabilized microtubules labeled with biotin and AZDye647, 30 μM unlabeled tubulin, 1.5 μM biotin-labeled tubulin and 1.5 μM AZDye647-labeled tubulin were mixed in BRB80 buffer supplemented with 1 mM GTP and incubated for 15 min at 37°C. Then, an equal amount of BRB80 supplemented with 40 μM taxol was added and further incubated for more than 15 min. The solution was loaded on BRB80 supplemented with 300 mM sucrose and 20 μM taxol and ultracentrifuged at 100,000 g for 5 min at 30°C. The pellet was resuspended in BRB80 supplemented with 20 μM taxol. Glass chambers were prepared by acid washing as previously described (9). Glass chambers were coated with PLL-PEG-biotin (SuSoS,

Dübendorf, Switzerland). Polymerized microtubules were flowed into streptavidin adsorbed flow chambers and allowed to adhere for 5–10 min. Unbound microtubules were washed away using assay buffer [90 mM Hepes pH 7.4, 50 mM $KCH_3COO$, 2 mM $Mg(CH_3COO)_2$, 1 mM EGTA, 10% glycerol, 0.1 mg/ml biotin–BSA, 0.2 mg/ml kappa-casein, 0.5% Pluronic F127, 2 mM ATP, and an oxygen scavenging system composed of PCA/PCD/Trolox. Purified motor protein was diluted to indicated concentrations in the assay buffer. Then, the solution was flowed into the glass chamber. An ECLIPSE Ti2-E microscope equipped with a CFI Apochromat TIRF 100XC Oil objective lens, an Andor iXion life 897 camera and a Ti2-LAPP illumination system (Nikon, Tokyo, Japan) was used to observe single molecule motility. NIS-Elements AR software ver. 5.2 (Nikon) was used to control the system.

***Microtubule gliding assays***

Heterodimers composed of KIF1A(1-393)::LZ::sfGFP::strep tag and KIF1A(1-393)::LZ::His tag were prepared as described in *Purification of heterodimers*. Tubulin was labeled with 5(6)-Carboxy tetramethylrhodamine (TMR) NHS ester (Fluoroprobes) and TMR labelled microtubules were prepared as described in *TIRF single-molecule motility assays*. Microtubule gliding assays were performed as described (10 , 11) with slight modifications. Glass chambers were coated with PLL-PEG-biotin (SuSoS). Streptavidin is adsorbed in flow chambers. Biotin-labelled anti-GFP antibodies diluted in BRB80 buffer were flowed into the chamber. Chamber was washed by assay buffer. Heterodimers composed of KIF1A(1-393)::LZ::sfGFP::strep tag and KIF1A(1-393)::LZ::His tag were diluted to indicated concentration in assay buffer and flowed into the chamber. Chamber was washed by assay buffer again. Microtubules diluted by assay buffer was flowed into the chamber and analyzed by the ECLIPSE Ti2-E microscope equipped with a CFI Apochromat TIRF 100XC Oil objective lens, an Andor iXion life 897 camera and a Ti2-LAPP illumination system (Nikon, Tokyo, Japan).

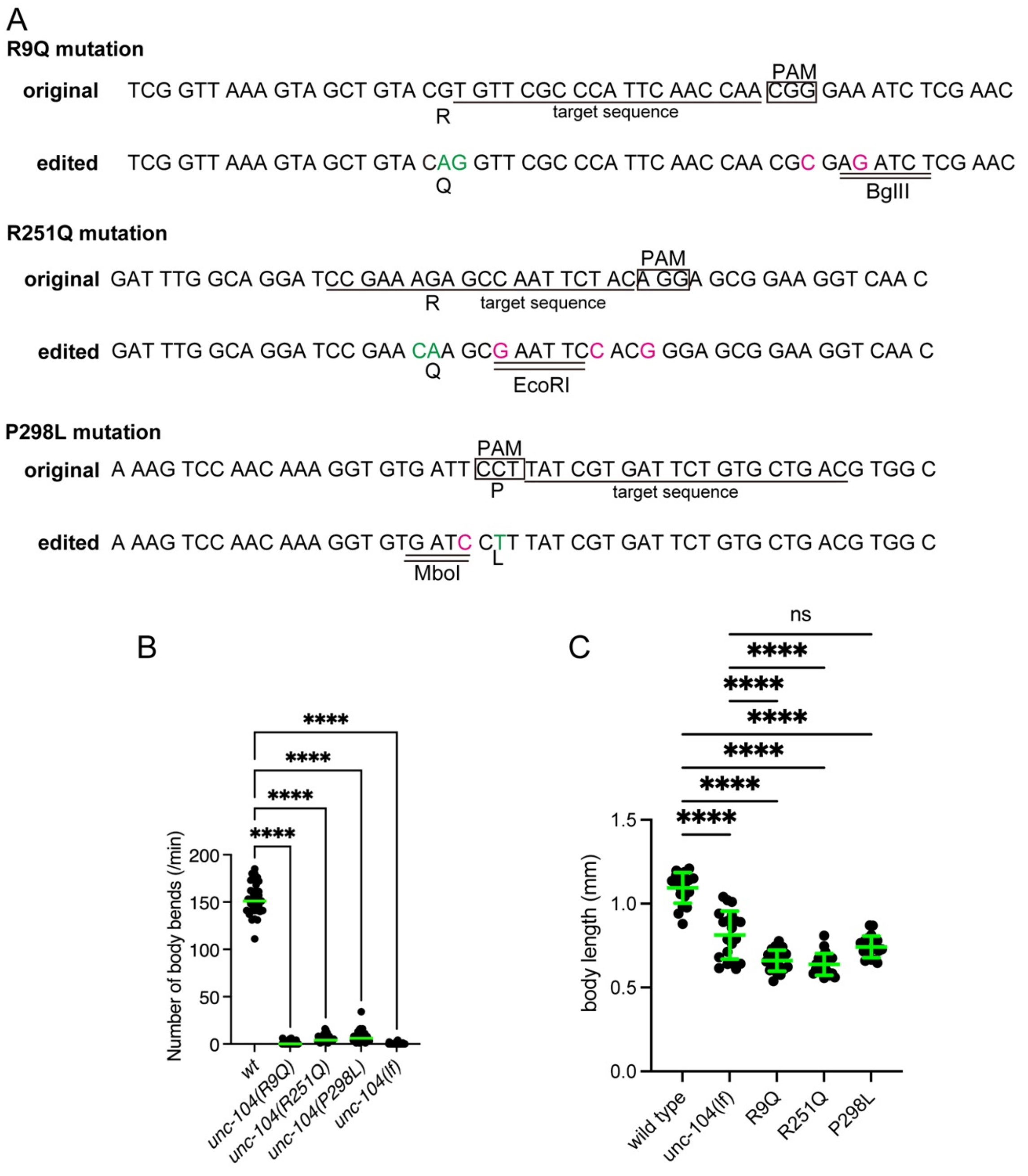


**Figure S1**

(A) Genome editing strategies. PAM (boxes) and target sequence (under lines) for cas9 is shown. Green font shows mutations to change amino acids. Magenta font shows silent mutations to disrupt target sequences and induce restriction enzyme sites. Double lines show restriction enzyme sites.

(B) Dot plots showing the result of swim test. The number of body bent in a water drop was counted for 1 min and plotted. Dots represent the number of bends from each worm. Green bars represent median value. Kruskal-Wallis test followed by Dunn's multiple comparison test. N = 20 worms for each genotype. ****, Adjusted P value < 0.0001.

(C) Dot plots showing the body length at 1-day adults. Dots represent the length from each worm. Green bars represent median value. Kruskal-Wallis test followed by Dunn's multiple comparison test. N = 20 worms for each genotype. ns, Adjusted P Value > 0.05 and statistically not significant. ****, Adjusted P value < 0.0001 compared to wt.

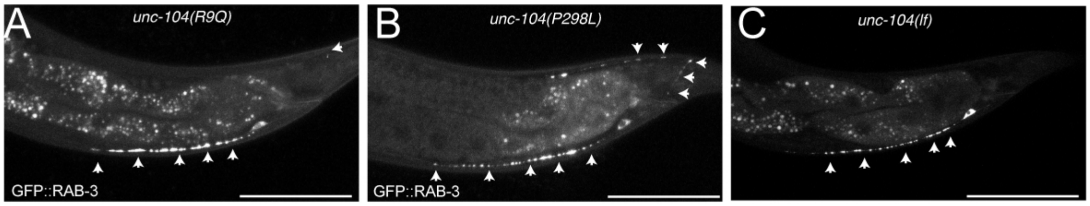


**Figure S2**
(A-C) Representative images showing the distribution of synaptic vesicles in DA9 neuron in *unc-104(R9Q)* (A), *unc-104(P298L)* (B) and *unc-104(lf)* (C). Synaptic vesicles are visualized by GFP::RAB-3. Arrow heads show mislocalization of synaptic vesicles in the dendrite and proximal axon. Bars, 50 µm.

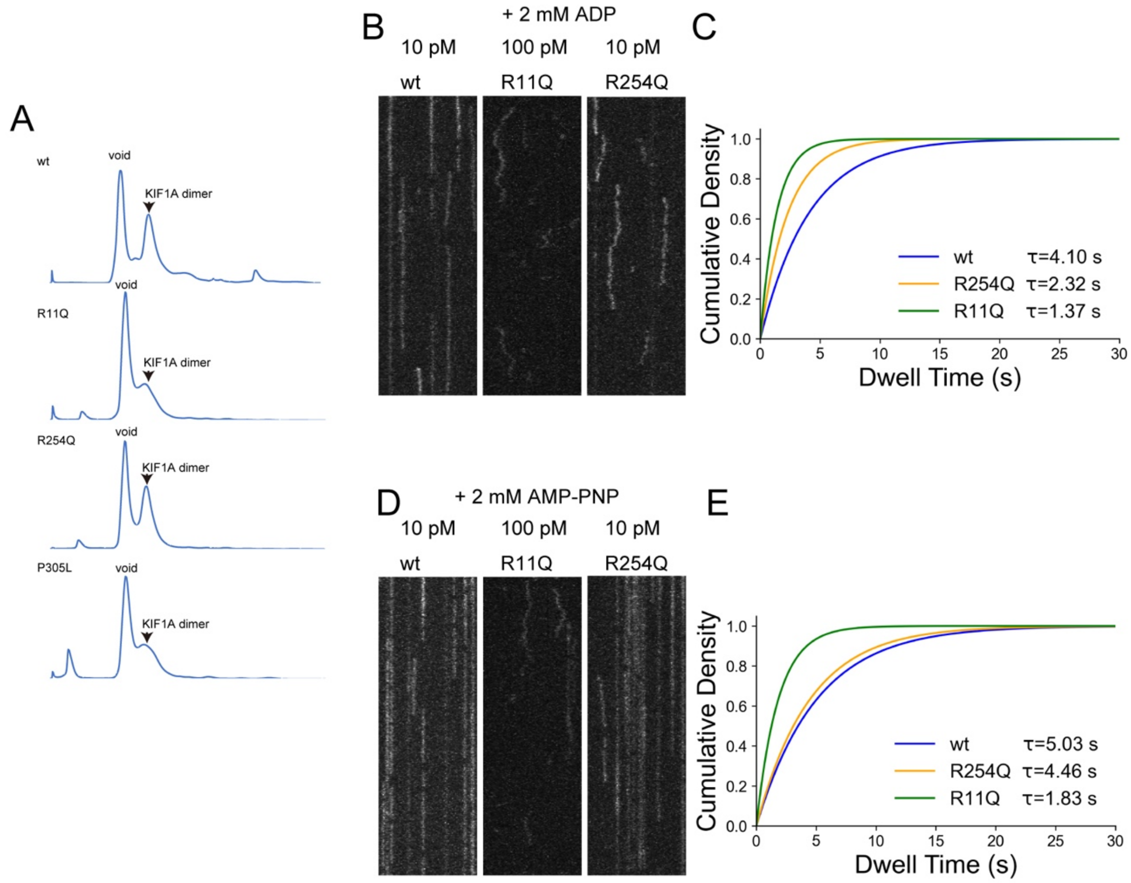


**Figure S3**

(A) Results of gel filtration. Recombinant KIF1A was recovered from the peak fraction shown by arrow heads.

(B and C) Binding of KIF1A with microtubules in the presence of ADP. (B) Representative kymographs showing the binding of 10 pM KIF1A(wt), 100 pM KIF1A(R11Q) and 10 pM KIF1A(R254Q). Vertical (30 seconds) and Horizontal (5 µm) windows are shown. (C) Cumulative density plots showing the dwell times of KIF1A(wt), KIF1A(R11Q) and KIF1A(R254Q) on microtubules in the presence of 2mM ADP. n = 169 (wt), 103 (R11Q) and 270 (R254Q) molecules.

(D and E) Binding of KIF1A with microtubules in the presence of an ATP analogue, AMP-PNP. (D) Representative kymographs showing the binding of 10 pM KIF1A(wt), 100 pM KIF1A(R11Q) and 10 pM KIF1A(R254Q). Vertical (30 seconds) and Horizontal (5 µm) windows are shown. (E) Cumulative density plots showing the dwell times of KIF1A(wt), KIF1A(R11Q) and KIF1A(R254Q) on microtubules in the presence of 2 mM AMP-PNP. n = 103 (wt), 115 (R11Q) and 140 (R254Q) molecules.

Note that molecules that did not bind or that did not dissociate in the observation time window were not included to the data. .

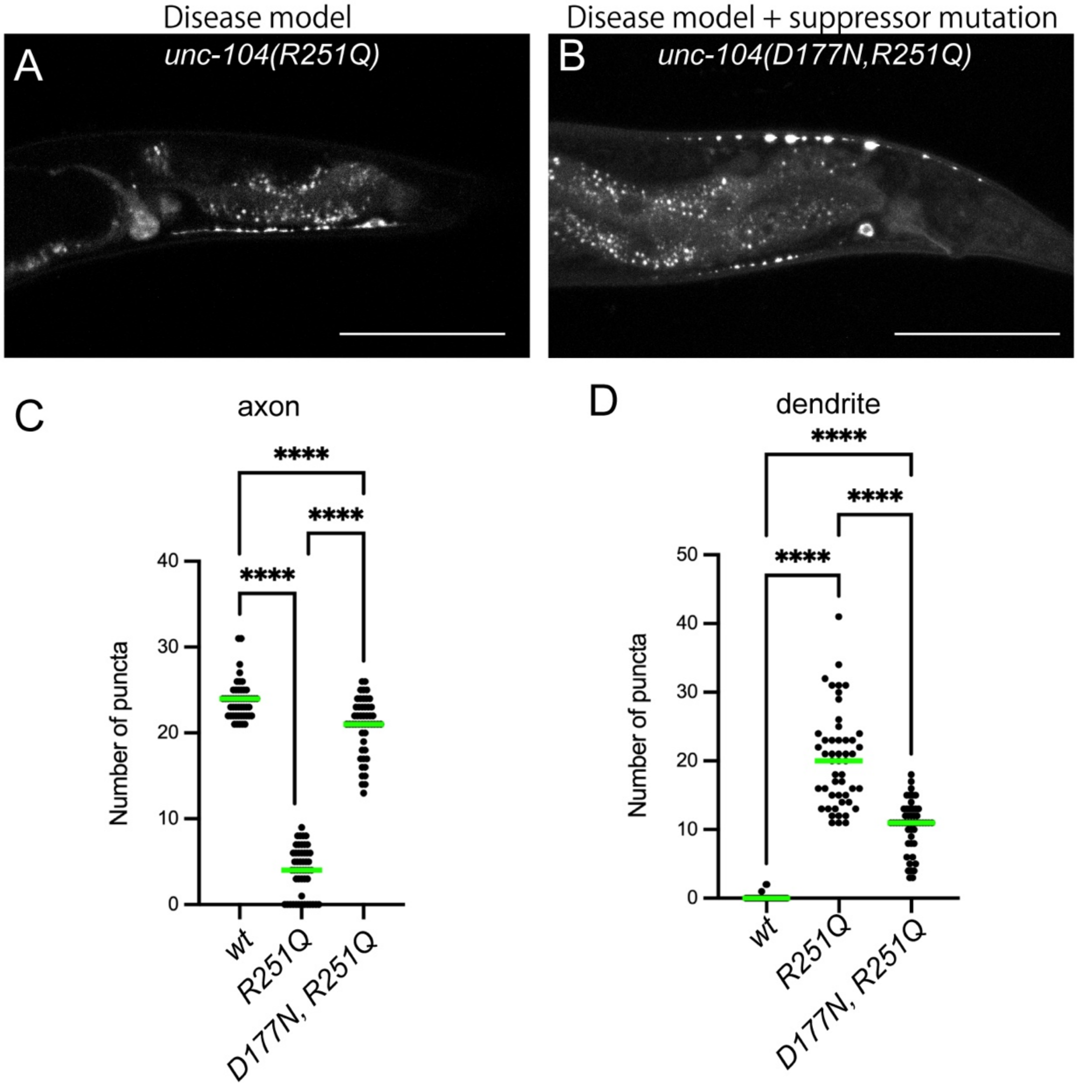


**Figure S4**

(A and B) Representative images showing the distribution of synaptic vesicles in DA9 neuron in *unc-104(R251Q)* (A), *unc-104(D177N, R251Q)* (B). Synaptic vesicles are visualized by GFP::RAB-3. Bars, 50 µm. Note that synaptic puncta are observed along the dorsal axon in the suppressor mutant.

(C and D) Dot plots showing the number of puncta in the dorsal axon (C) and ventral dendrite (D) of DA9 neuron. Green bars represent median value. N = 50 worms from each genotype. Kruskal-Wallis test followed by Dunn's multiple comparison test. ****, Adjusted P value < 0.0001.

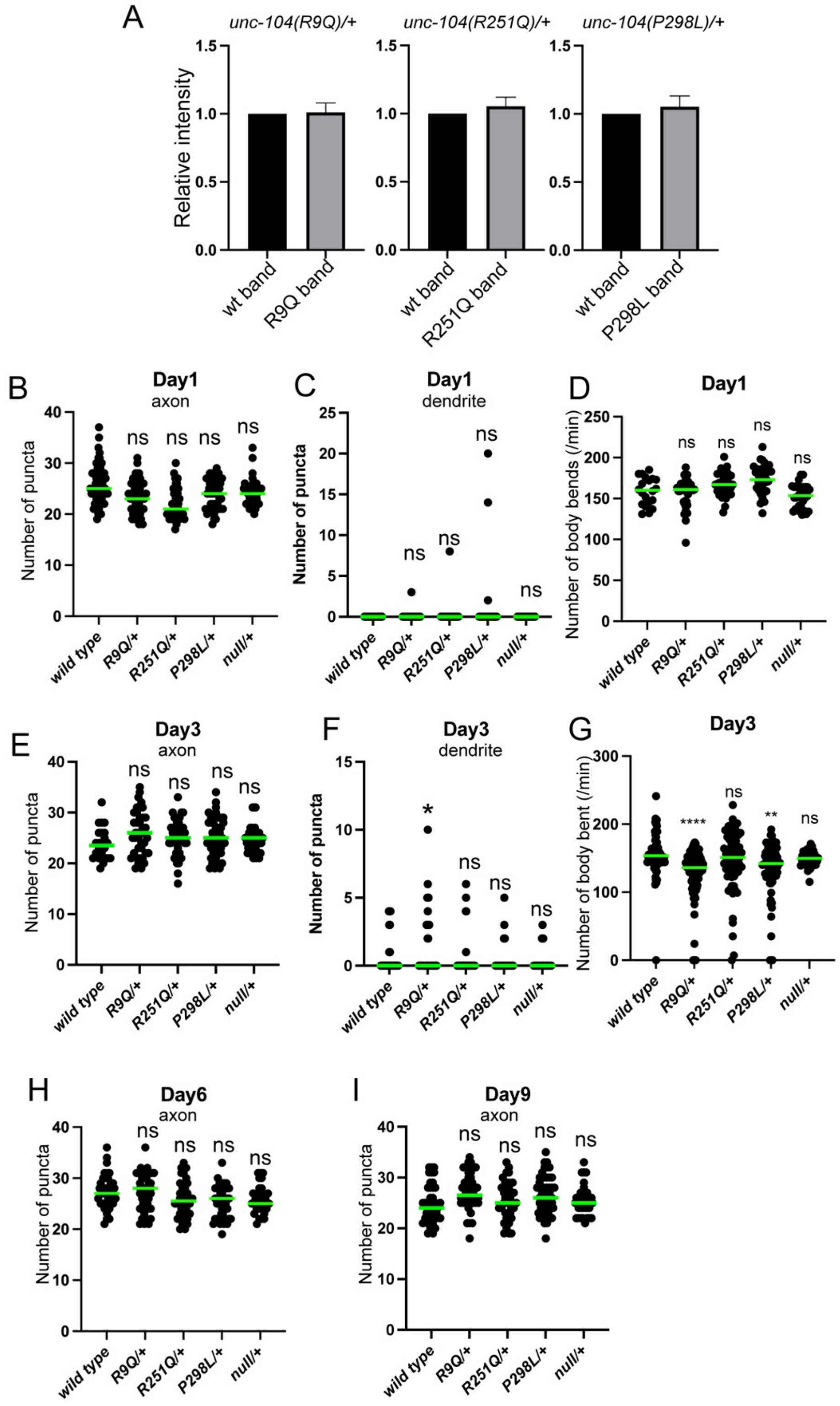


**Figure S5**

(A) Bar graphs showing the relative expression level of wild type and mutant mRNA in heterozygotes. After RT-PCR was performed, PCR products were digested by restriction

enzyme sites (Figure S1A) to discriminate wild type and mutant copies. Agarose gel electrophoresis followed by ethidium bromide staining was performed. Band intensities were quantified and adjusted by the size. Bars represents the relative intensities. Error bars show S.D.. N = 3 independent experiments.
(B and C) Dot plots showing the number of puncta in the dorsal axon (B) and ventral dendrite (C) of DA9 neuron at day 1. Green bars represent median value. Kruskal-Wallis test followed by Dunn's multiple comparison test. N = 60 (wt), 46 (R9Q/+), 43 (R251Q/+), 51 (P298L/+) and 47 (null/+) worms. ns, Adjusted P Value > 0.05 and statistically not significant compared to wt.
(D) Dot plots showing the result of swim test at 1 day. The number of body bent in a water drop was counted for 1 min and plotted. Dots represents the number of bends from each worm. Green bars represent median value. Kruskal-Wallis test followed by Dunn's multiple comparison test. N = 76 (wt), 87 (R9Q/+), 74 (R251Q/+), 65 (P298L/+) and 39 (null/+) worms. ns, Adjusted P Value > 0.05 and statistically not significant compared to wt.
(E and F) Dot plots showing the number of puncta in the dorsal axon (E) and ventral dendrite (F) of DA9 neuron at day 3. Green bars represent median value. N = 24 (wt), 40 (R9Q/+), 40 (R251Q/+), 45 (P298L/+) and 45 (null/+) worms. Kruskal-Wallis test followed by Dunn's multiple comparison test. ns, Adjusted P Value > 0.05 and statistically not significant. *, Adjusted P Value < 0.05. Compared to wt.
(G) Dot plots showing the result of swim test at day 3. The number of body bent in a water drop was counted for 1 min and plotted. Dots represents the number of bends from each worm. Green bars represent median value. N = 80 (wt), 94 (R9Q/+), 62 (R251Q/+), 72 (P298L/+) and 38 (null/+) worms. Kruskal-Wallis test followed by Dunn's multiple comparison test. ns, Adjusted P Value > 0.05 and statistically not significant. **, Adjusted P Value < 0.01, ****, Adjusted P Value < 0.0001. Compared to wt.
(H and I) Dot plots showing the number of puncta in the dorsal axon of DA9 neuron at 6 days (H) and 9 days (I). Green bars represent median value. Kruskal-Wallis test followed by Dunn's multiple comparison test. (H) N = 39 (wt), 43 (R9Q/+), 36 (R251Q/+), 46 (P298L/+) and 32 (null/+) worms. (I) N = 39 (wt), 40 (R9Q/+), 37 (R251Q/+), 49 (P298L/+) and 37 (null/+) worms. ns, Adjusted P Value > 0.05 and statistically not significant compared to wt control.

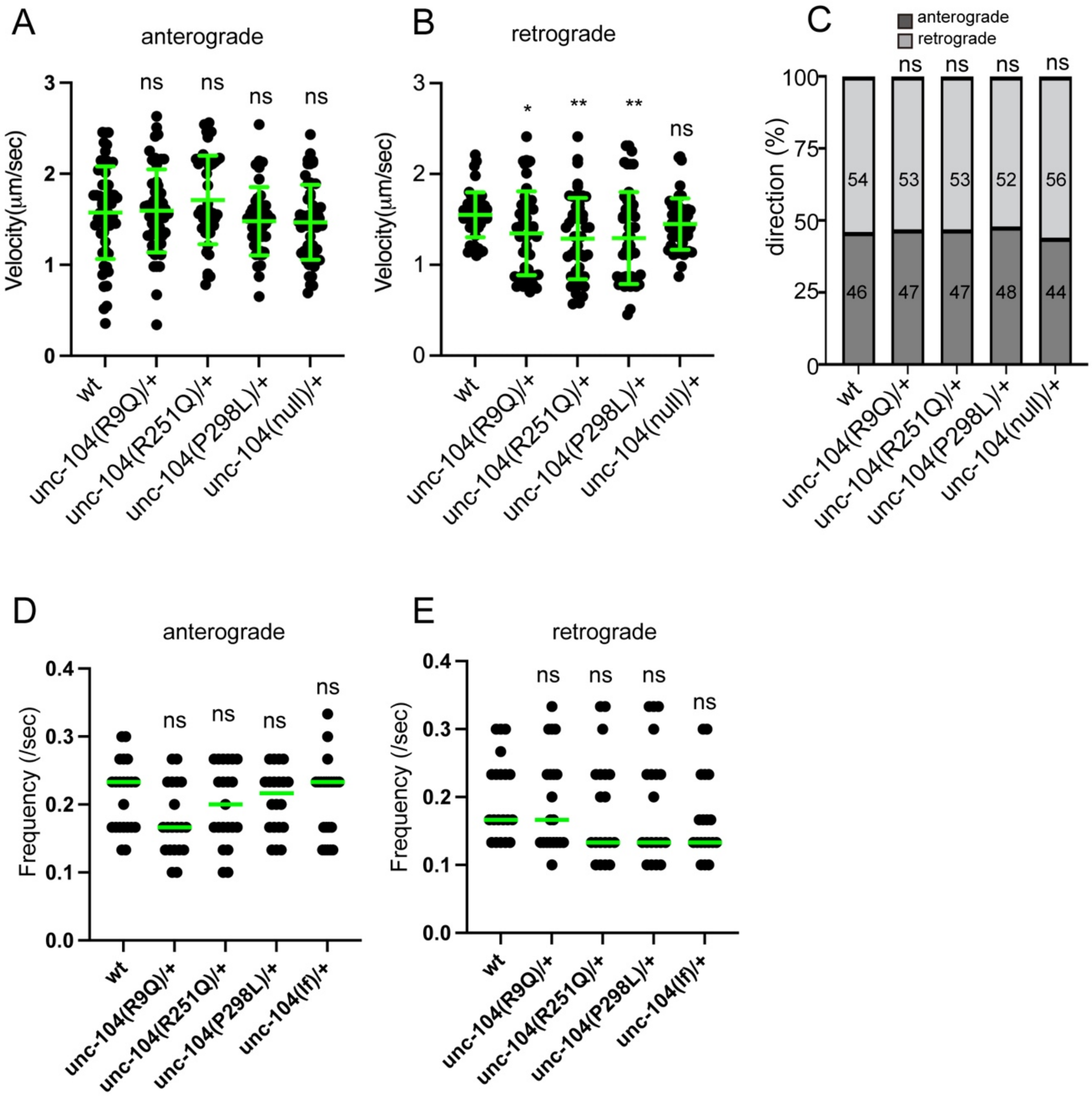


**Figure S6**
(A and B) The velocity of dendritic transport. The velocity of anterograde transport (i.e. transport from the cell body to dendritic tip) (A) and retrograde transport (i.e. transport from the dendritic tip to cell body) (B) are shown as dot plots. Green bars show mean ± S.D.. (A) n = 56 (wild type), 56 (R9Q/+), 45 (R251Q/+), 42 (P298L/+) and 56 (null/+) vesicles from at least 10 independent worms. (B) n = 50 vesicles from each genotype from at least 10 independent worms. Kruskal-Wallis test followed by Dunn's multiple comparison test. ns, Adjusted P Value > 0.05 and no significant statistical difference. *, Adjusted P Value < 0.05. **, Adjusted P Value < 0.01. Compared to wt worms.
(C) Directionality of vesicle movement. The number in the bar graph shows the actual percentage. Chi-square test. ns, Adjusted P Value > 0.05 and statistically not significant compared to wt worms.
(D and E) Frequency of dendritic transport. The frequency of anterograde transport (D) and retrograde transport (E) are shown as dot plots. Each dot represent data from each

worm. Green bars represent median value. (D) N = 22 (wt), 20 (R9Q/+), 21 (R251Q/+), 20 (P298L/+) and 22 (null/+) independent worms. (E) N = 20 (wt), 20 (R9Q/+), 19 (R251Q/+), 19 (P298L/+) and 19 (null/+) independent worms. Kruskal-Wallis test followed by Dunn's multiple comparison test. ns, Adjusted P Value > 0.05 and no significant statistical difference compared to wt worms.

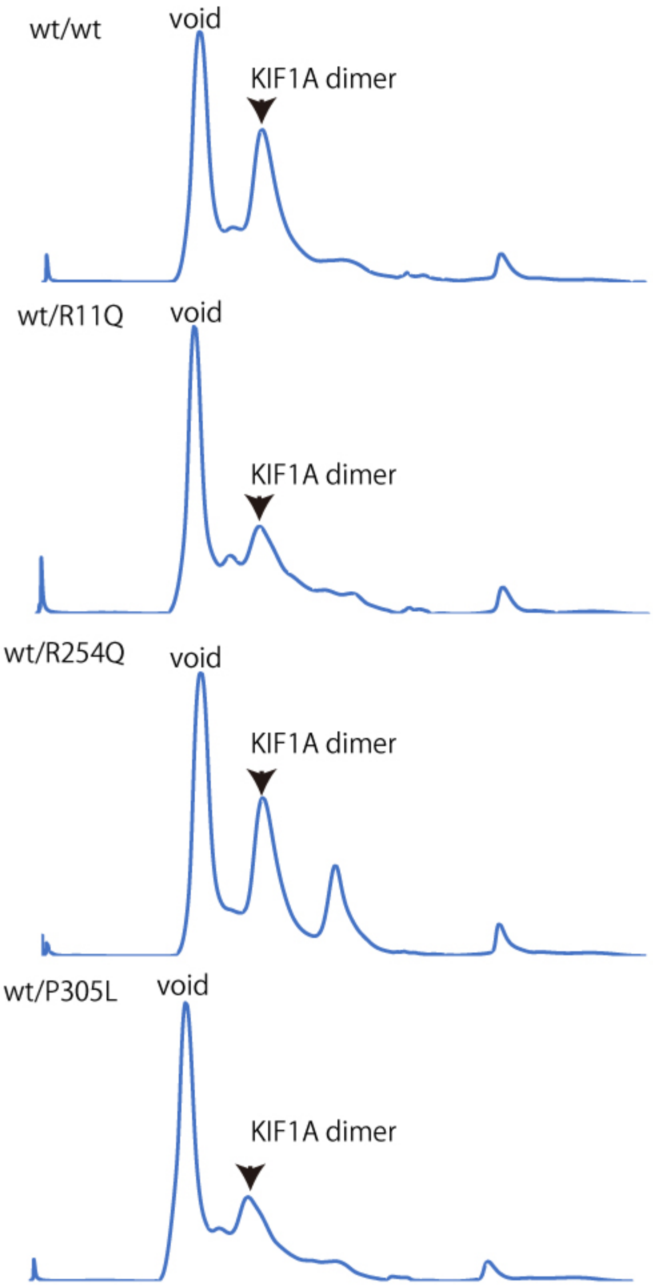


**Figure S7**
Results of gel filtration. Recombinant KIF1A was recovered from the peak fraction shown by arrow heads.

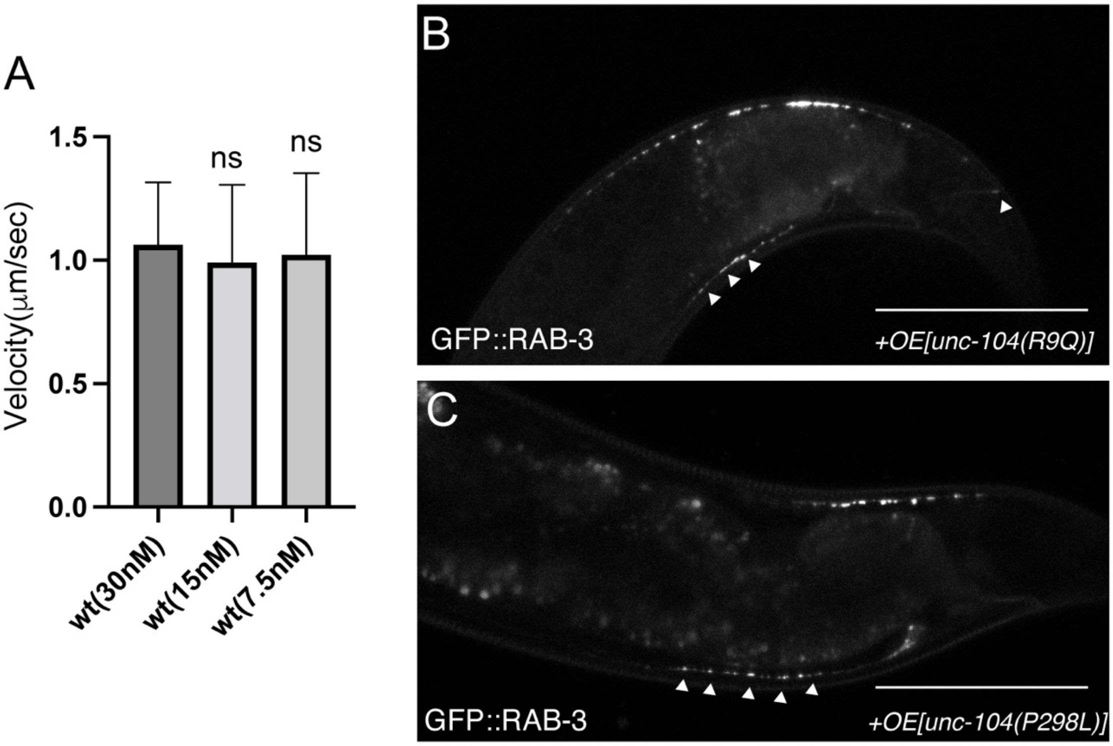


**Figure S8**

(A) Microtubule gliding assays at different concentrations. Bars and error bars represent mean and SD, respectively. N = 146 (30 nM wt homodimers), 112 (15 nM wt homodimers), 20 (7.5 nM wt homodimers), microtubules from at least three independent experiments. Kruskal-Wallis test followed by Dunn's multiple comparison test. ns, Adjusted P Value > 0.05 and no significant statistical difference compared to 30 nM. . (B and C) Representative images showing the localization of synaptic vesicles in UNC-104(R9Q)-expressing worm (B) and UNC-104(P298L)-expressing worm (C). Arrow heads show synaptic-vesicle accumulated puncta that are mislocalized in the proximal axon and dendrite. Bars, 50 μm.

### Supplementary Table S1 DNA sequences for genome editing

| mutation + restriction enzyme | sgRNA expression plasmid (name, target sequence) | repair template (sequence) |
|---|---|---|
| unc-104(R9Q) + BglII | unc-104(V6M)#1, 5'- TGTTCGCCCATTCAACCAA -3' | 5'- TTTCAGATGTCATCGGTTAAAGTAGCTGTACagGTTCGCCCATTCAACCAACGcGAgATCTCGAACACTTCAAAATGTGTCCTTCAAGTAAATGGAAAT -3' |
| unc-104(R251Q) + EcoRI | unc-104(A252V)#4, 5'- CCAATTCTACAGGAGCGGA -3' | 5'-ACTGAGAAGCATTCAAAAATTTCTTTGGTTGATTTGGCAGGATCCGAAcaAGCgAATTCcACgGGAGCGGAgGGTCAACGACTAAAAGAAGGAGCAAATA -3' |
| unc-104(P298L) + MboI | unc-104(P298L), 5'- GTCAGCACAGAATCACGATA -3' | 5'- AGTTTAATTTATTTATAGTCAACAAAGAAGAAAAAGTCCAACAAAGGTGTGATcCgTTATCGTGATTCTGTGCTGACGTGGCTTCTGAGAGAAAATCT -3' |
| ben-1 | ben-1#1, 5'- AGTGATATCCGATGAGCAT -3' | |

#### Supplementary Table S2 Strain list

| strain | genotype | description |
|---|---|---|
| N2 | wild type | Obtained from CGC |
| TV1229 | *wyIs85[Pitr-1::gfp::rab-3]* | Figure 1 and 2, Obtained from Dr. Kang Shen (Stanford University) and crossed with N2 |
| OTL150 | *unc-104(e1265): wyIs85* | *unc-104(e1265)* was obtained from CGC. *unc-104(e1265)* is *unc-104(lf)*. |
| TV9315 | *wyIs251[Pmig-13::gfp::rab-3]* | Figure 1 and 2, Obtained from Dr. Kang Shen (Stanford University) |
| OTL129 | *unc-104(jpn23); wyIs85* | Figure 1 and 2. UNC-104(R251Q) mutation. KIF1A(R254Q) model |
| OTL130 | *unc-104(jpn29);wyIs85* | Figure 1 and 2. UNC-104(P298L) mutation. KIF1A(P305L) model |
| OTL131 | *unc-104(jpn30);wyIs85* | Figure1 and 2. UNC-104(R11Q) mutation. KIF1A(R11Q) model |
| OTL132 | *unc-104(jpn23); wyIs251* | Figure 2 and 6. UNC-104(R251Q) mutation. KIF1A(R254Q) model |
| OTL133 | *unc-104(jpn29); wyIs251* | Figure 6. UNC-104(P298L) mutation. KIF1A(P305L) model |
| OTL134 | *unc-104(jpn30); wyIs251* | Figure 6. UNC-104(P298L) mutation. KIF1A(P305L) model |
| OTL140 | *unc-104(jpn23jpn44); wyIs85* | unc-104(R251Q. D177N) mutation. Figure 4. Data from this strain is shown in this paper. |
| OTL141 | *unc-104(jpn23jpn45); wyIs85* | unc-104(R251Q. D177N) mutation. |
| OTL142 | *wyIs85; jpnEx102[Pmig-13::unc-104]* | Figure 8 |
| OTL143 | *wyIs85; jpnEx103[Pmig-13::unc-104]* | Figure 8 |
| OTL144 | *wyIs85; jpnEx104[Pmig-13::unc-104(R9Q)]* | Figure 8 |
| OTL145 | *wyIs85; jpnEx105[Pmig-13::unc-104(R9Q)]* | Figure 8 |
| OTL146 | *wyIs85; jpnEx106[Pmig-13::unc-104(R251Q)]* | Figure 8 |
| OTL147 | *wyIs85; jpnEx107[Pmig-13::unc-104(R251Q)]* | Figure 8 |
| OTL148 | *wyIs85; jpnEx108[Pmig-13::unc-104(P298L)]* | Figure 8 |
| OTL149 | *wyIs85; jpnEx109[Pmig-13::unc-104(P298L)]* | Figure 8 |
| OTL151 | *unc-104(jpn23)/mIn1; wyIs85* | Figure 5 and 6 |
| OTL152 | *unc-104(jpn29)/mn1; wyIs85* | Figure 5 and 6 |
| OTL153 | *unc-104(jpn30)/mIn1; wyIs85* | Figure 5 and 6 |
| OTL154 | *unc-104(tm819)/mIn1; wyIs85* | Figure 5 and 6. unc-104(tm819)/mIn1 was obtained from Mitani lab. *unc-104(tm819)* is *unc-104(null)* |

**Supplementary Table S3  Plasmid list**

| Plasmid | Description | Backbone |
|---|---|---|
| pSN643 | KIF1A(1-393)::LZ::mScarlet-I::Strep tag | pET28a |
| pSN656 | KIF1A(1-393)(R11Q)::LZ::mScarlet-I::Strep tag | pET28a |
| pSN657 | KIF1A(1-393)(R254Q)::LZ::mScarlet-I::Strep tag | pET28a |
| pSN658 | KIF1A(1-393)(P305L)::LZ::mScarlet-I::Strep tag | pET28a |
| pSN672 | KIF1A(1-393)::LZ::His tag | pET21a |
| pSN685 | KIF1A(1-393)(R11Q)::LZ::His tag | pET21a |
| pSN686 | KIF1A(1-393)(R254Q)::LZ::His tag | pET21a |
| pSN687 | KIF1A(1-393)(P305L)::LZ::His tag | pET21a |

1.
